# Growth-dependent sensory bet-hedging enhances collective navigation

**DOI:** 10.64898/2026.02.13.705748

**Authors:** Fotios Avgidis, Ruta Jurgelyte, Thierry Emonet, Thomas S. Shimizu

**Affiliations:** Center for Living Systems, AMOLF Institute, Amsterdam, 1098 XG, The Netherlands; Department of Molecular, Cellular, and Developmental Biology, Yale University, New Haven, 06511, CT, USA; Quantitative Biology Institute, Yale University, New Haven, 06511, CT, USA; Department of Physics, Yale University, New Haven, 06511, CT, USA

## Abstract

Phenotypic heterogeneity in microbes can be a double-edged sword — boosting individual survival in uncertain environments, but potentially compromising beneficial population coordination. Dissecting how microbes resolve this tradeoff is challenging because interactions span multiple scales—from molecular interactions to single-cell behavior, population dynamics, and environmental feedback. Here, we address this question for the *Escherichia coli* chemotaxis system, which implements both individual motile explorations and collective resource exploitations using the same cellular machinery. We quantify the heterogeneity of individual sensory and behavioral phenotypes, as well as the abundance of key signaling proteins during growth in various environments, and test their impact on population-scale collective migration. We identify growth rate as a key environment-dependent physiological variable governing not only the mean but also the variance of sensory phenotypes. Remarkably, rather than hindering population coordination, we find that sensory diversity benefits collective chemotactic navigation in uncertain environments. Strong heterogeneity of expressed phenotypes enhances readiness to multiple sensory cues, and the required population coordination is achieved by phenotypic filtering of that diversity by the collective behavior itself. These results reveal a sensory bet-hedging strategy for collective navigation during population growth: diversity in sensitivity to nutrients currently being consumed is reduced to promote focused exploitation, while it is increased for nutrients not yet encountered to enhance exploration for new growth opportunities.

## 1 Introduction

The survival of an organism depends upon its capacity to adeptly perceive and respond to environmental cues, such as nutrient resources required for growth. This faculty is particularly important for microbes such as bacteria, which inhabit diverse ecosystems characterized by rapid and unpredictable changes, such as terrestrial soils or the mammalian gut [1]. Growing populations are often confronted with exhaustion of a particular nutrient source, as well as new encounters with alternative resources. A strategy to cope with such unpredictability is to establish a population that mixes individuals of different phenotypes. This diversification, often called “bet-hedging”, can enhance collective readiness of populations to a range of environmental changes, by maintaining subpopulations of individuals prepared for contrasting future scenarios [2–8].

Yet phenotypic diversity also poses challenges for population-level coordination. Many collective microbial processes such as quorum sensing [9, 10], biofilm formation [11, 12], and antibiotic defense [13, 14] depend on secreted signals or public goods reaching some threshold, and can fail if the fraction of non-secreting (or “cheater”) cells is too high. Microbial developmental programs such as sporulation waves in B. subtilis [15, 16] and fruiting body formation in M. xanthus [17, 18], as well as host manipulation and invasion strategies by pathogens [19, 20] tend to require synchronized state transitions across the population. And in chemotactic bacteria, efficient collective exploitation of nutrient resources requires traveling-wave range expansion by populations with matched chemotactic velocities [21–23]. In all of these cases, nonzero diversity tends to confer bet-hedging advantages (cells that do not invest in the collective phenotype often are endowed with reproductive or survival advantages), but excessive phenotypic heterogeneity clearly becomes detrimental.

What are the strategies and mechanisms by which microbial populations negotiate this tension? In this study, we address this question in the *Escherichia coli* chemotaxis system, which controls motile navigation both at the level of individual cells by biasing a run-and-tumble random walk, and at the level of the population, by generating chemotactic population waves that migrate at a constant speed. Previous work established that the chemosensory system controlling this motile behavior exhibits phenotypic diversity suggestive of bet-hedging [24–30], and that it can rapidly modulate its phenotype distribution without changes in gene expression through a diversity-tuning mechanism based on covalent modification of allosterically interacting chemoreceptors [8, 31]. However, that posttranslational mechanism works by effectively filtering the underlying diversity in protein counts, which can itself vary on a slower timescale due to stochastic gene expression. We therefore began by characterizing single-cell sensory phenotypes and stochastic gene expression under various environmental conditions, before examining their implications for collective navigation performance.

At the single-cell level, *E. coli* moves through space by a run-and-tumble random walk [32]. To bias this random walk towards attractants and away from repellents, *E. coli* makes use of the chemotaxis pathway, one of the best characterized models of biological signaling. *E. coli* cells detect chemical and physical signals through arrays of transmembrane receptors composed of five different types of chemoreceptors. The two primary receptors, Tar (which mainly senses the amino acid aspartate) and Tsr (which mainly senses the amino acid serine), make up 90% of the total chemoreceptor population when *E. coli* cells are grown in rich media. The other three chemoreceptors—Tap, Trg, and Aer—are present at substantially lower copy numbers [33]. The receptors modulate the activity of the kinase CheA, which phosphorylates the response regulator CheY, producing CheY-P. Binding of attractant molecules to chemoreceptors reduces the activity of CheA, leading to a decrease in the intracellular level of CheY-P due to the activity of the phosphatase CheZ. CheY-P interacts with the flagellar motor, reducing the bacterium’s tendency to tumble. Consequently, attractant binding increases the duration of the bacterium’s runs, biasing the random walk in the desired direction.

We characterized the diversity of *E. coli* ‘s sensory and motility phenotypes, as well as the expression of its two major chemoreceptors, Tar and Tsr, under various growth conditions. Specifically, by combining the single-cell FRET microscopy method developed in our lab [27] with a microfluidic device that allows tracking hundreds of swimming trajectories simultaneously [34] and fluorescent fusions of Tar and Tsr, we quantified sensory and behavioral diversity alongside gene expression noise in isogenic bacteria under nutrient-rich and nutrient-poor environments. Our findings reveal that changes in nutrient composition and cell density alter not only the average sensory phenotype but also the degree of cell-to-cell variability in sensory responses. In contrast, motility phenotypes—specifically the tumble bias and run speed, which together define the run-and-tumble random walk in the absence of a chemical gradient—remain constant across all tested environments.

By combining experiments and mathematical analysis of a multi-species allosteric (MWC) model [35, 36], we demonstrate that the standing level of sensory diversity can be explained by changes in just one cellular random variable governed by stochastic gene expression: the Tar/Tsr receptor protein ratio.

By utilizing custom microfluidic chemostats amenable to single-cell fluorescence microscopy (variants of the “mother machine” [37]), we found that the chemical composition of the growth environment, rather than growth duration, dictates the distribution of the Tar/Tsr ratio through a growth rate-dependent gene expression mechanism.

Finally, by generating populations of cells with increased diversity in sensory phenotypes, we show that a high level of standing diversity confers a discernible advantage for chemotactic navigation under behavioral contexts where populations are challenged with different chemoattractants as environmental sensory cues. This advantage arises from the rapid filtering for the best-performing phenotypes by the collectively migrating chemotactic population wave, and occurs independently of gene expression changes —– a novel population-level adaptation mechanism we recently predicted and verified experimentally [34, 38].

The gene-expression-based diversity adaptation mechanism we describe here operates on a timescale of hours, acting in parallel with the faster, non-genetic collective migration adaptation mechanism that occurs within minutes, and the posttranslational diversity tuning mechanism which operates on the sensory adaptation timescale of ∼ 10 seconds.

## 2 Results

### 2.1 Single-cell FRET reveals substantial diversity in ligand sensitivity across an isogenic population of cells

To probe the degree of cell-to-cell variation in ligand sensing within a genetically identical population, we performed dose-response measurements by stimulating *E. coli* cells with pulses of the attractant amino acid L-serine that binds the major chemoreceptor Tsr, while monitoring the output of the signaling pathway in single cells using an *in vivo* CheYZ FRET sensor [27] for the activity of the kinase CheA (Figure 1a). *E. coli* has a sensory adaptation system consisting of the enzymes CheR and CheB that modulate the sensitivity of the chemoreceptors via covalent modifications that maintain the kinase output of the pathway to a constant steady-state level. To decouple diversity in ligand sensitivity from diversity in receptor modification induced by the two adaptation enzymes, we deleted the genes encoding CheR and CheB. We performed dose-response measurements (see methods) with cells grown as batch culture in nutrient-rich (TB) medium containing all 20 amino acids until mid-exponential phase, corresponding to an optical density (OD) of 0.45. The resulting dose-response data were fitted with a sigmoidal Hill function of the form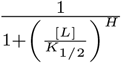, where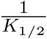, represents ligand sensitivity and H the cooperativity between receptors.

**Fig. 1.**
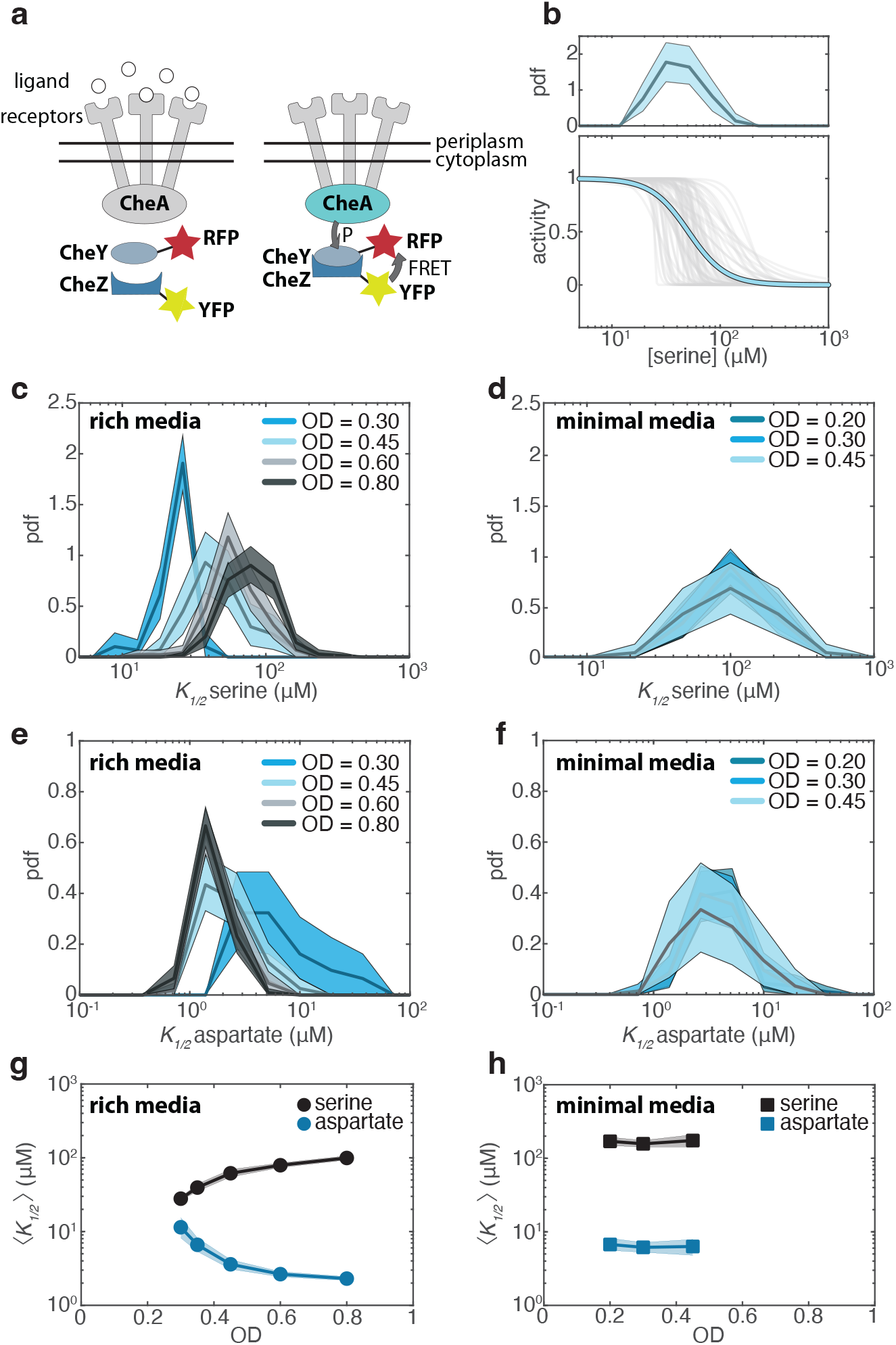
Single-cell FRET reveals extensive environment-dependent diversity in ligand sensitivity across an isogenic bacterial population. **(a)** Schematic of the *in vivo* FRET assay. The kinase CheA phosphorylates the response regulator CheY producing CheY-P in the absence of ligand binding to receptors. CheZ dephosphorylates CheY. The FRET signal between fluorescently labeled CheZ and CheY is proportional to the CheA activity. **(b)** (Bottom) single-cell (gray) and population-averaged (blue) Hill function fits to L-serine dose-response data obtained from a single experiment using adaptation-deficient cells grown to OD = 0.45 in rich media. (Top) Histogram of the half-maximal concentration *K*_1*/*2_, extracted from the single-cell dose response curves shown above. **(c)** Histograms of L-serine *K*_1*/*2_ values obtained from single-cell dose-response curves for cells grown to different ODs in rich media. Each distribution represents data from a single experiment. Shaded areas indicate 95% confidence intervals obtained via bootstrap resampling of the data. **(d)**Same as (c), but for cells grown in minimal media **(e)** Histograms of L-aspartate *K*_1*/*2_ values obtained from single-cell dose-response curves for cells grown to different ODs in rich media. Each distribution represents data from a single experiment. Shaded areas indicate 95% confidence intervals obtained via bootstrap resampling. Figure legend and color scheme are consistent with panel (c). **(f)** As in (e), but for cells grown in minimal media. Figure legend and color scheme are consistent with panel (d). **(g)** Scaling of mean L-serine and L-aspartate *K*_1*/*2_ as a function of OD for cells grown in rich media. Shaded areas indicate 95% confidence intervals obtained via bootstrap resampling. **(h)** Same as (g), but for cells grown in minimal media.

This analysis revealed that single-cell dose-response curves can differ substantially from the population-average curve (Figure 1b). Extracting the *K*_1*/*2_ parameter from these single-cell curves, which corresponds to the ligand concentration necessary to reduce the kinase activity by 50%, revealed a substantial degree of diversity, with the *K*_1*/*2_ values of individual cells spanning approximately an order of magnitude (Figure 1b), consistent with our previous findings [8, 27, 31] .

### 2.2 Sensitivity distributions shift as a function of cell density in rich media, but not in minimal media

Considering that *E. coli* can be found in a variety of nutrient-diverse ecological niches, we wondered how chemosensory diversity is influenced by the growth environment. We cultured isogenic populations of cells in the same nutrient-rich batch culture and suspended growth by harvesting at four different cell densities, ranging from OD = 0.30 to OD = 0.80, where cells experience different nutrient concentrations [39–41]. Serine dose-response experiments revealed that the distribution of the half-maximal concentration *K*_1*/*2_, inversely related to sensitivity, shifts towards higher values with increasing cell density (Figure 1c). Notably, the diversity in *K*_1*/*2_, measured as the coefficient of variation (CV, defined as standard deviation divided by the mean), increases considerably with cell density, ranging from CV = 0.20 at OD = 0.30 to CV = 0.45 at OD = 0.80.

We then examined how diversity varies as a function of cell density in an environment devoid of chemotactic cues. Cells were grown in minimal media without amino acids, using glycerol as a carbon source, which does not elicit a chemotactic response [42]. Compared to cells grown in rich media, these cells exhibited reduced sensitivity (higher *K*_1*/*2_) in their responses to serine (Figure 1d). Notably, in minimal media, the distribution of *K*_1*/*2_ remained independent of cell density, in contrast to the density-dependent shifts observed in rich media. Furthermore, the coefficient of variation of *K*_1*/*2_ remained very high (CV ∼ 0.50) across all cell densities in minimal media.

Finally, we wondered how the sensitivity to the other major *E. coli* chemoattractant, L-aspartate, which binds the chemoreceptor Tar, is modulated by the growth environment. We performed experiments identical to those described above, substituting serine with aspartate. Interestingly, in rich media, we observed the opposite trend compared to serine: cells exhibited higher sensitivity to aspartate (lower *K*_1*/*2_) at higher cell densities (Figure 1e). However, as with serine, the distribution of aspartate *K*_1*/*2_ was independent of cell density in minimal media (Figure 1f). We found that the coefficient of variation of aspartate *K*_1*/*2_ was also notably high, ranging from CV = 0.91 at OD = 0.30 to CV = 0.34 at OD = 0.80. In minimal media, the CV of aspartate *K*_1*/*2_ remained very high (CV ∼ 0.80) across all cell densities.

In summary, these results reveal an inverse scaling of serine and aspartate sensitivity with cell density in rich media: as cell density increases, sensitivity to aspartate increases (lower *K*_1*/*2_), whereas sensitivity to serine decreases (higher *K*_1*/*2_) (Figure 1g). In contrast, in minimal media, the sensitivities to both attractants are typically lower than in rich media and remain invariant with respect to cell density (Figure 1h).

### 2.3 Tar/Tsr ratio distribution shifts as a function of cell density in rich media, but not in minimal media

Next, we sought to identify the source of this modulation of chemotactic diversity. Previous studies have shown that stochastic gene expression can lead to significant variability in protein levels among isogenic cells, with one study suggesting that proteins in *E. coli* have a baseline variation of 30% in their expression level [43]. Of the dozen or so protein species in the chemotaxis network of *E. coli*, we focused our attention on two, the aspartate receptor Tar and the serine receptor Tsr, since prior studies have demonstrated that the relative ratio of these receptors significantly impact chemotactic behavior, influencing factors such as preference for aspartate/methyl-aspartate or serine [44–46], the switch between cryophilic and thermophilic preference [47, 48], and the switch between base-seeking and acid-seeking behavior [49].

To quantify cell-to-cell variability in the Tar/Tsr ratio, we engineered functional fluorescent fusions of the *tar* and *tsr* genes at their native chromosomal loci, fusing *tar* to mCherry and *tsr* to a monomeric form of YFP (Figure 2a) and quantified their single-cell expression levels by fluorescence microscopy (see methods). Using the same parent strain of *E. coli*, MG1655, and identical growth conditions as in our FRET experiments, we measured Tar/Tsr ratio variation under the same conditions in which we assessed chemotactic diversity. When grown in nutrient-rich media to mid-exponential phase (OD = 0.45), cells exhibited substantial cell-to-cell variation in the Tar/Tsr ratio (Figure 2b), with a CV of 0.35. This value is comparable to, though slightly lower than, the Tar/Tsr ratio CV reported by a study that used flow cytometry measurements in a different *E. coli* strain, RP437, which exhibited a CV of 0.45 at OD = 0.51 [47]. The Tar/Tsr ratio CV predicted for the same RP437 strain using single-cell FRET data was 0.5 at OD = 0.45 [27]. These differences are likely attributable to expression differences across *E. coli* strains, which are known to be often substantial even among close relatives [33].

**Fig. 2.**
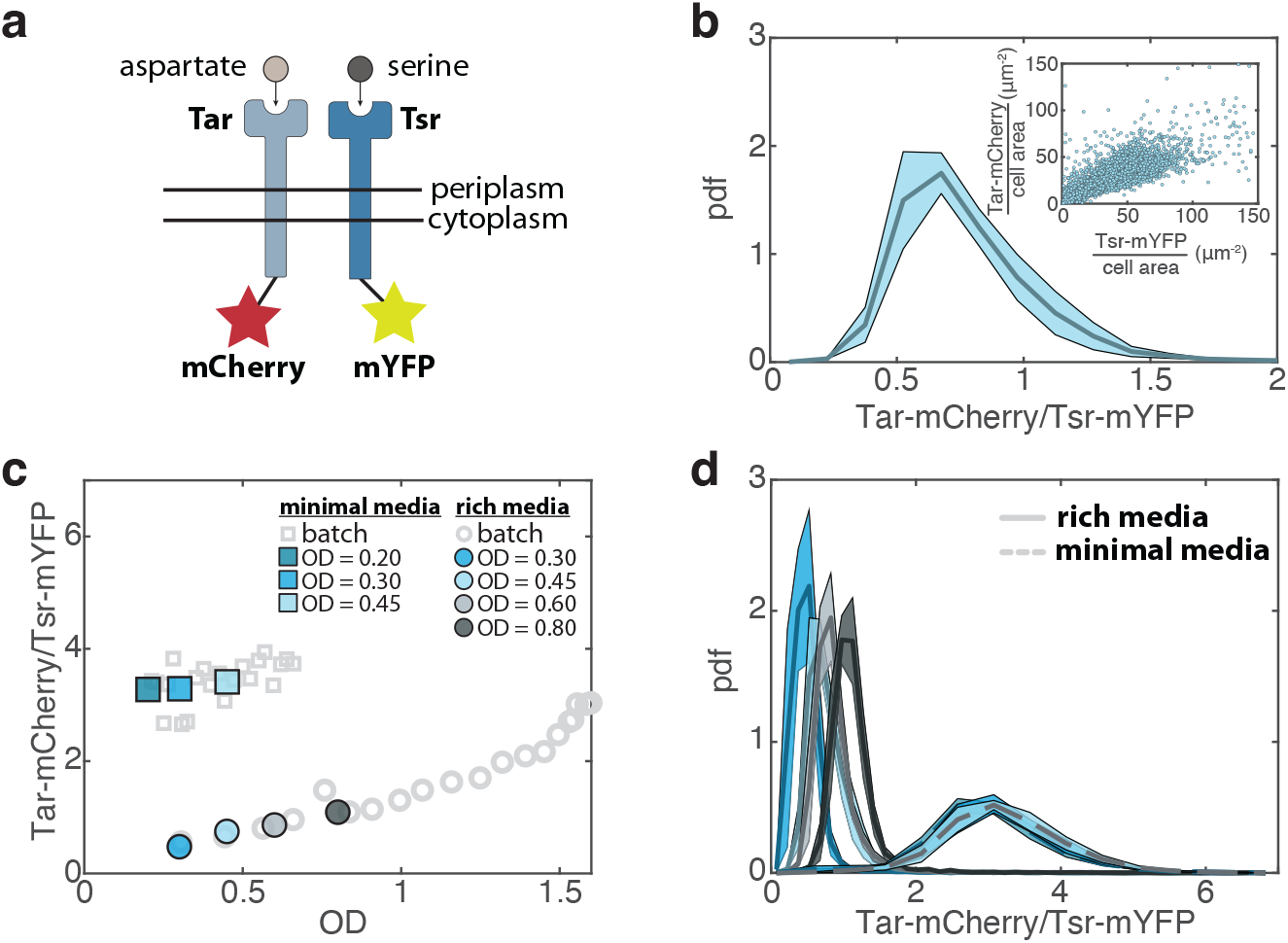
Fluorescent fusions of major chemoreceptors reveal extensive environment-dependent diversity in chemoreceptor composition across an isogenic bacterial population. **(a)** Schematic of the receptor protein quantification assay. The chemoreceptor Tar, which binds the amino acid L-aspartate, is fused at its native chromosomal locus to the fluorophore mCherry. The chemoreceptor Tsr, which binds the amino acid L-serine, is fused at its native chromosomal locus to the fluorophore mYFP. Imaging parameters were calibrated using a two-color fluorescent repressor-operator system (FROS) standard to equate the measured mCherry/YFP fluorescent intensity ratio to the true Tar/Tsr receptor abundance ratio (EDF 3). **(b)** Histogram of the Tar/Tsr chemoreceptor ratio measured with fluorescence microscopy for cells grown to OD = 0.45 in rich media, growth condition identical to the one used in figure 1b. The solid line represents the mean of four independent biological replicates, while shaded areas indicate the standard error of the mean across replicates. Inset: Correlation between Tar-mCherry and Tsr-mYFP expression levels. Each point represents a single cell, derived from four independent biological replicates. Tar and Tsr expression levels are strongly correlated (*R*^2^ = 0.70). **(c)** Relationship between the Tar/Tsr chemore-ceptor ratio and OD for cells grown in rich (circles) and minimal (squares) media. Gray hollow points represent the mean of 40 technical replicates measured using a plate reader. Colored points represent the mean Tar/Tsr ratio measured via fluorescence microscopy from four (rich media) or three (minimal media) independent biological replicates for each OD. **(d)** Histograms of the Tar/Tsr chemoreceptor ratio measured via fluorescence microscopy for cells grown to different ODs in rich (solid line) or minimal (dashed line) media. Lines represent the mean distribution of all independent biological replicates, while shaded areas indicate the standard error of the mean across replicates. The color scheme is consistent with panel c.

Next, we assessed how the Tar/Tsr ratio evolves as a function of cell density. Using a plate reader, we continuously monitored Tar and Tsr expression and the OD of the cell culture. Cells initially exhibited a low Tar/Tsr ratio, which increased as cell density rose, consistent with previous findings (Figure 2c) [44, 46–48]. Notably, single-cell fluorescence microscopy revealed that the CV of the Tar/Tsr ratio decreased with increasing cell density, ranging from a CV of 0.43 at OD = 0.30 to a CV of 0.23 at OD = 0.80 (Figure 2d), consistent with modeling predictions of a previous study [27]. This trend mirrors that of the aspartate *K*_1*/*2_, whose CV decreases with increasing cell density, and contrasts with the trends observed for serine *K*_1*/*2_, whose CV increases with cell density. The increase in the Tar/Tsr ratio at higher optical densities was primarily driven by increased Tar expression, while Tsr levels remained mostly constant (Figure EDF 4). We also observed a strong correlation between Tar and Tsr expression at all optical densities (Figure 2b, inset and EDF 5), which is expected since the genes encoding these receptors—though located in separate operons—are regulated by a common transcriptional regulator, FliA, along with all other chemotaxis genes [47, 50]. Their expression is also likely influenced by global cellular factors that affect the expression of the whole proteome, such as ribosome and polymerase availability [51, 52].

To further investigate how the receptor ratio evolves in different environments, we repeated our plate reader assay in minimal media. In minimal media, the Tar/Tsr ratio remained mostly constant as the cell density increased (Figure 2c) and the distribution of Tar/Tsr ratios was identical across all sampled optical densities (Figure 2d). This observation aligns with the invariant *K*_1*/*2_ distributions observed as a function of OD in minimal media (Figures 1d and 1f). The average Tar/Tsr ratio in minimal media was skewed towards higher Tar levels (Tar/Tsr ∼ 3.3; figure 2d), similar to the Tar/Tsr ratio measured at high optical densities in rich media (Tar/Tsr = 3.25 at OD ∼ 1.6; figure 2c). Additionally, the CV of the Tar/Tsr ratio was lower in minimal media compared to rich media, with a CV of approximately 0.31, across all optical densities. For both rich and minimal media, the standard deviation in Tar and Tsr protein copy numbers increased with increasing mean copy numbers (EDF 5c), a result consistent with a previous study that characterized the expression of 1018 proteins in *E. coli* using YFP fusions [43].

Our findings suggest that the shift toward a higher Tar fraction at elevated optical densities in rich media (Figure 2c) is the primary driver of increased aspartate sensitivity under these conditions (Figure 1g). Although the total Tsr expression remained approximately constant (EDF 4b), the sharp decrease in serine sensitivity at higher cell densities (Figure 1g) is likely a consequence of chemoreceptor cluster organization. Tar and Tsr are known to form mixed clusters within the cell membrane [53]; thus, as Tar expression rises with cell density, newly synthesized Tar receptors dilute pre-existing clusters composed primarily of Tsr molecules. This dilution effect reduces overall serine sensitivity despite the approximately constant Tsr expression levels.

### 2.4 Diversity in the Tar/Tsr ratio is sufficient to quantitatively predict chemosensory diversity

To quantitatively compare the measured diversity of *K*_1*/*2_ to the measured diversity in the Tar/Tsr ratio, we turned to theoretical modeling. We employed a mixed-species Monod-Wyman-Changeux (MWC) model [35], which has been successful in fitting a number of *in vivo E. coli* chemotaxis single-cell FRET data [8, 27, 31, 36]. Using this model, we generated a serine dose-response curve for each cell, by fixing the total number of MWC receptors (*N*_*total*_) and using as a single model input their measured Tar/Tsr ratio (see methods). Then, we extracted the single-cell *K*_1*/*2_ from each dose-response curve using the same hill function we used to fit experimental FRET data. When comparing the CV of *K*_1*/*2_ obtained from our FRET data to the CV of *K*_1*/*2_ predicted through the MWC model from our receptor ratio data, we found excellent quantitative agreement across all experimental conditions (Figure 3a). This result alone indicates that a significant portion, if not all, of the observed sensory diversity in the chemotaxis system of *E. coli* can be attributed to the stochastic gene expression of its major chemoreceptors, Tar and Tsr. To further establish direct causality between receptor expression diversity and *K*_1*/*2_ diversity, we generated a population of cells with artificially high levels of receptor ratio variability. To achieve that, we cloned the same Tar-mCherry fusion we engineered on the chromosome of *E. coli* into an inducible vector under the control of a leaky Lac promoter. By transforming the receptor-labeled strain with this vector and tuning its induction level in conjunction with tuning the harvesting OD of the strain (which affects receptor ratio variability in rich media; Figure 2d), we were able to generate a population of cells with approximately the same mean Tar/Tsr ratio as wild-type cells harvested at OD = 0.80 in rich media, but with an approximately two-fold greater Tar/Tsr ratio CV (Figure 3b). We then transformed a strain amenable to FRET experiments with a vector carrying the wild-type Tar receptor under the control of the same leaky Lac promoter. This generated a population of cells with approximately the same mean *K*_1*/*2_ as wild-type cells harvested at OD = 0.80 in rich media, but with an approximately three-fold higher *K*_1*/*2_ CV (Figure 3c). Importantly, MWC modeling again revealed excellent quantitative agreement between measured *K*_1*/*2_ CV and *K*_1*/*2_ CV predicted by the model calibrated by receptor-ratio measurements (Figure 3c, inset), thus demonstrating direct causality between receptor expression diversity and diversity in ligand sensitivity.

**Fig. 3.**
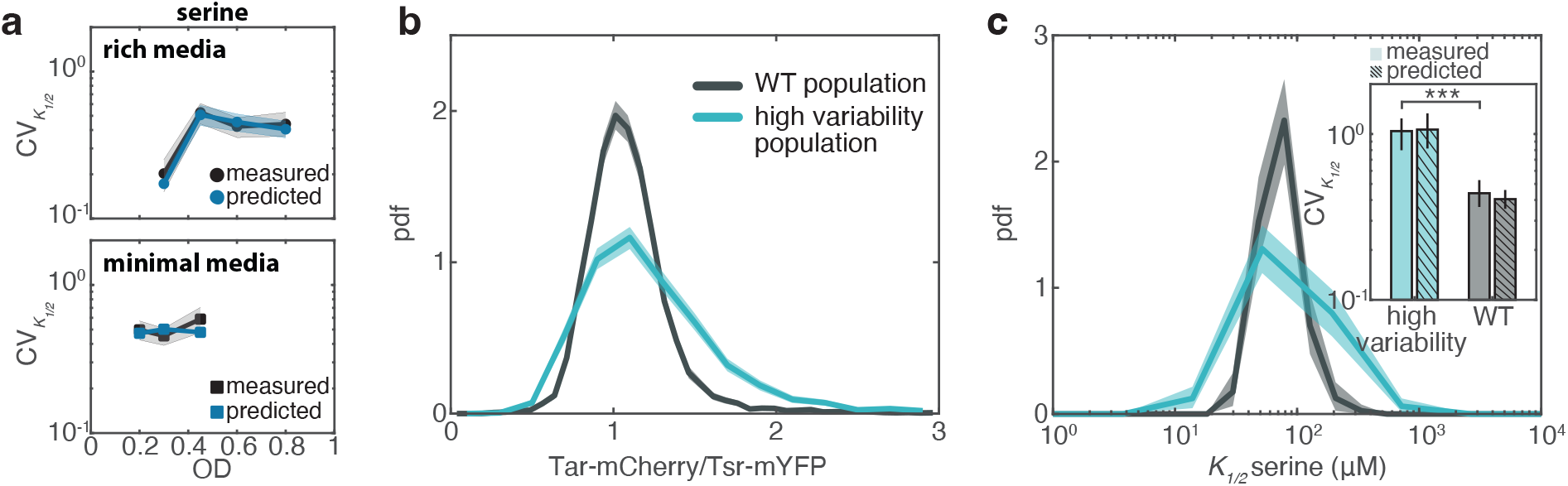
Prediction of chemosensory diversity from Tar/Tsr ratio variation, and establishing causality. **(a)** (Top) Black points represent the measured coefficient of variation (CV) of the the half-maximal concentration *K*_1*/*2_ as a function of OD in rich media, obtained from the single-cell dose-response experiments in Figure 1c. Blue points represent the predicted CV of *K*_1*/*2_ as a function of OD in rich media, derived through the mixed-species MWC model using single-cell receptor ratio data from Figure 2d as an input. Shaded areas indicate 95% confidence intervals obtained through bootstrap resampling. (Bottom) Same as the top panel, but for cells grown in minimal media. The CV was either measured from the single-cell dose-response experiments in Figure 1d (black points) or predicted from the mixed-species MWC model using single-cell receptor ratio data from Figure 2d as input (blue points). The total number of MWC receptors (*N*_*total*_) was fixed to 32 for all rich media experiments and 100 for all minimal media experiments. **(b)** Histograms of the Tar/Tsr chemoreceptor ratio measured via fluorescence microscopy for wild-type cells (WT population; replotted OD = 0.80 data from Figure 2d) and for a population with artificially increased Tar/Tsr ratio variability (high variability population). Solid lines represent the histograms of all Tar/Tsr ratios from four independent biological replicates (WT population) or a single experiment (high variability population). Shaded areas indicate 95% confidence intervals obtained through bootstrap resampling. **(c)** Histograms of L-serine *K*_1*/*2_ values obtained from single-cell dose-response curves for wild-type cells (WT population; replotted OD = 0.80 data from figure 1c) or cells with artificially increased Tar/Tsr ratio variability (high variability population). Solid lines represent the histograms from a single experiment, with shaded areas indicating 95% confidence intervals obtained through bootstrap resampling. Inset: Measured CV of *K*_1*/*2_ for WT and high variability populations (solid-colored bars), along with predicted CV of *K*_1*/*2_ obtained from the mixed-species MWC model using the receptor ratio distributions in Figure 3c as input (dashed-colored bars) and a fixed total number of MWC receptors (*N*_*total*_ = 32). Error bars indicate 95% confidence intervals obtained through bootstrap resampling. The means are statistically significant different, as determined by a two-sample t-test (p = 2.9x10^*−*12^).

### 2.5 Growth environment, not growth time, dictates the Tar/Tsr ratio distribution

We then sought to understand the cause of the observed shift in the Tar/Tsr ratio as a function of cell density in rich media. To test whether the total growth time determines the receptor ratio, we diluted a saturated bacterial culture into fresh rich media at three different final concentrations (Figure 4a). We then harvested all cultures at the same optical density (OD = 0.45). As expected, increasing the dilution factor by two-fold extended the total growth time by one division cycle (Figure 4b, inset). However, the resulting receptor ratio distributions were identical across all conditions (Figure 4b), demonstrating that growth time has no effect on receptor expression. This result is consistent with a previous study that determined the mean receptor ratio at different growth times using quantitative immunoblotting [44].

**Fig. 4.**
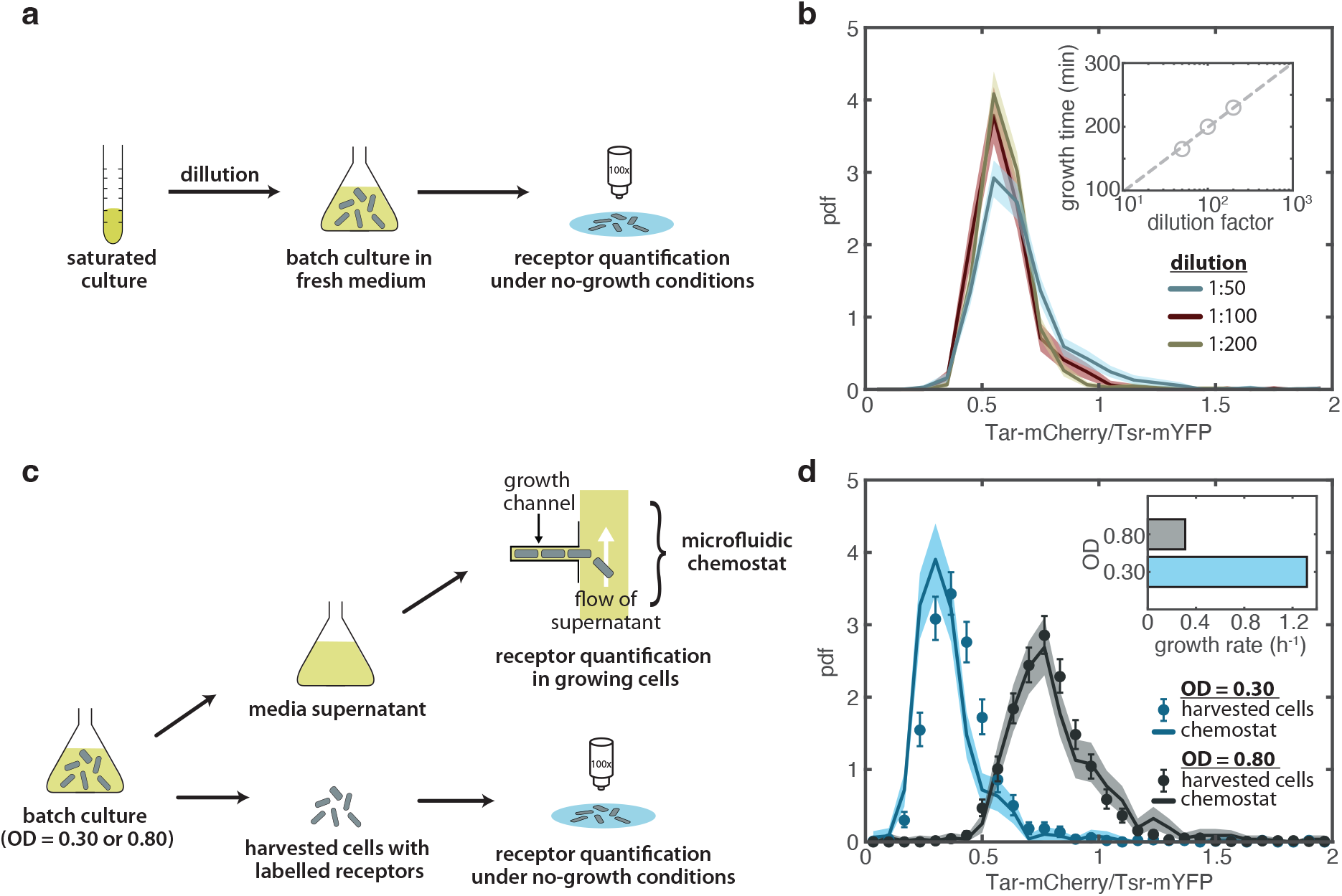
Growth environment, rather than growth time, determines Tar/Tsr ratio through growth rate-dependent chemoreceptor expression. **(a)** Schematic of the experimental protocol to assess the effect of growth time on the chemoreceptor ratio distributions. Cells from the same saturated culture were diluted to three different concentrations in fresh rich media and all cultures were harvested at the same optical density (OD = 0.45). **(b)** Histograms of the Tar/Tsr chemoreceptor ratio, measured via fluorescence microscopy, for the three different dilutions in a single experiment. The receptor ratio distribution is independent of the dilution factor. Error bars indicate 95% confidence intervals obtained through bootstrap resampling. Inset: Growth time as a function of dilution. **(c)** Schematic of experimental protocol to assess the effect of the growth environment on chemoreceptor ratio distributions. Cells were grown in rich media to OD = 0.30 and OD = 0.80, after which they were separated from their supernatant. The Tar/Tsr chemoreceptor ratio was measured via fluorescence microscopy in both the harvested cells and in cells growing inside a microfluidic chemostat at steady state, where they were exposed to the collected supernatant. **(d)**Histograms of the Tar/Tsr chemoreceptor ratio for harvested cells (solid lines) and for cells growing inside the chemostat while exposed to supernatant (points). Solid lines and points represent the histograms of all measured Tar/Tsr ratios from a single experiment. Shaded areas and error bars indicate 95% confidence intervals obtained through bootstrap resampling. Inset: growth rate of the cells in the batch culture at OD = 0.30 and OD = 0.80, assessed using a plate reader. Bar plots are the means of 40 technical replicates.

To explore whether the chemical environment determines the Tar/Tsr ratio instead, we employed a custom microfluidic chemostat amenable to single-cell microscopy (a “mother machine” [37], see methods). Within the microfluidic chemostat, the growth medium is constantly replenished, and excess cells are washed away, maintaining a constant environment and steady-state growth conditions. We grew cells in rich media to the two extremes of the cell densities examined earlier (OD = 0.30 and OD = 0.80) and measured their receptor ratios. In parallel, we collected the supernatant from both cultures. We then exposed cells growing inside the microfluidic chemostat to the collected supernatant and, after a few hours of growth in each supernatant, we determined their receptor ratios (Figure 4c). Strikingly, we found excellent quantitative agreement between the receptor ratio distributions of the harvested cells and those of cells growing at steady state in the corresponding supernatant (Figure 4d). We also found that growth rate decreases at higher ODs (Figure 4d, inset).

Taken together, these results demonstrate that the receptor ratio of *E. coli* does not follow a predetermined developmental trajectory, nor does it employ a timer-based mechanism that resets upon entry into a new environment. Instead, cells sense changes in their environment and dynamically adjust their receptor ratio—and, by extension, their chemotactic behavior—accordingly.

### 2.6 Growth rate governs sensory bet-hedging through chemoreceptor ratio modulation

Next, we investigated the molecular mechanism underlying the environment-dependent regulation of the Tar/Tsr ratio. As shown above, cells at low optical density (OD) grow substantially faster than those at high OD (Figure 4d, inset), prompting us to ask whether the Tar/Tsr ratio regulation is mediated by growth rate. Indeed, in rich media, growth rate decreases as a function of cell density, slowing as the culture approaches saturation (Figure 5a, inset). In minimal media, however, growth rate remains approximately constant during exponential phase but can be modulated by supplementing the medium with different carbon sources. To examine how growth rate affects the receptor protein concentrations, we measured Tar protein concentrations across a range of growth rates in both rich and minimal media using a plate reader assay (see methods). We observed an approximately a 10-fold decrease in Tar concentration as the growth rate increased across all environments (Figure 5a, top), consistent with previous studies [54]. This inverse relationship is likely driven by catabolite repression mediated by cyclic adenosine monophosphate (cAMP) and the cAMP receptor protein (CRP) [54]. In contrast, Tsr concentration remained constant regardless of growth rate (Figure 5a, bottom). The Tar and Tsr chemoreceptor concentrations obtained from batch, plate reader, data closely matched those obtained via single-cell fluorescence microscopy (Figure 5a), supporting the robustness of these measurements.

**Fig. 5.**
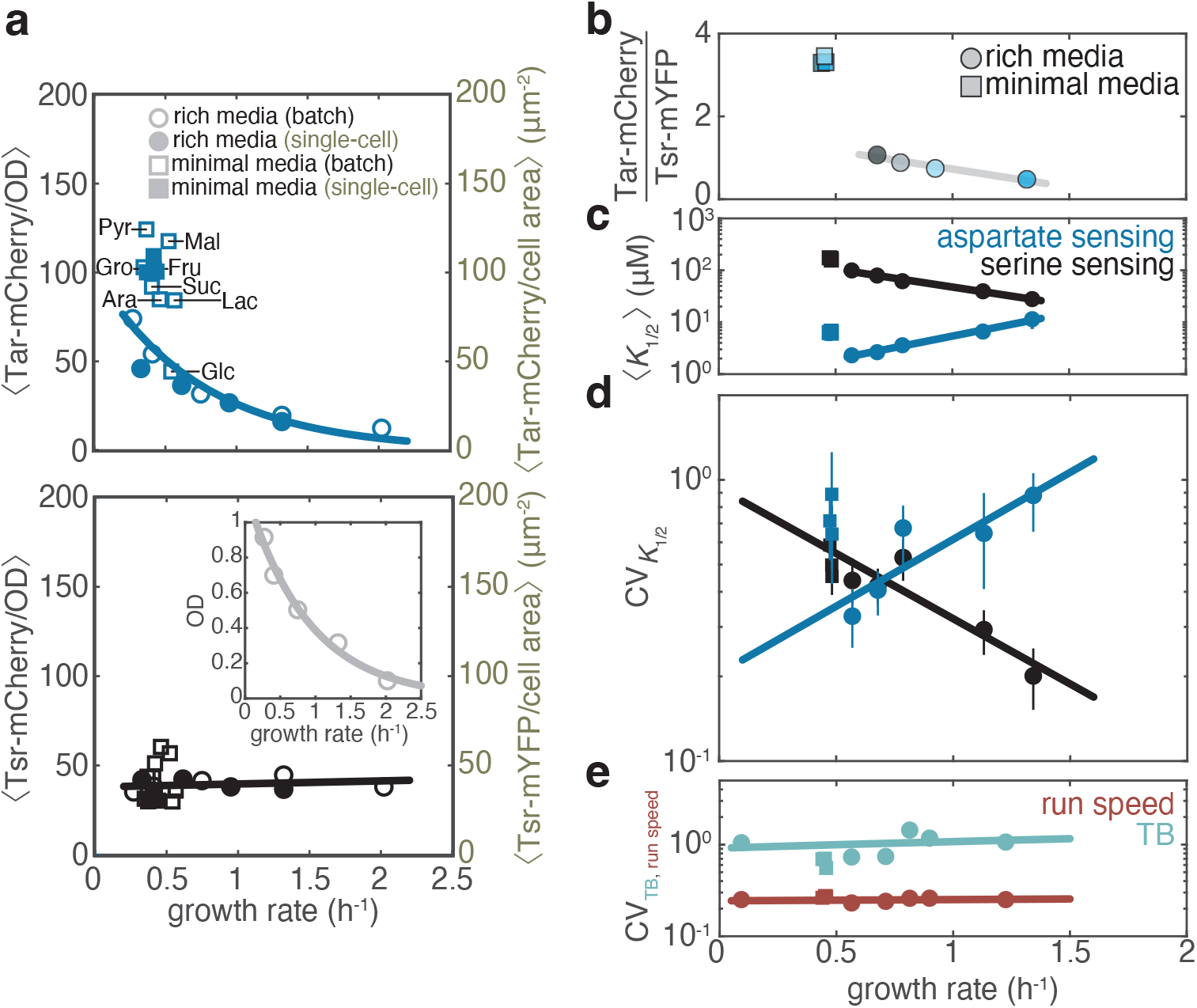
Bacteria regulate their sensory diversity based on growth rate. **(a)** (Top) Concentration of Tar proteins as a function of growth rate. Protein concentration in batch cultures (hollow points) is defined as the mean fluorescence between two time points during growth divided by the mean biomass (OD) at the same two time points. Carbon sources used to supplement minimal media are indicated as: Gro: 0.5% v/v glycerol; Pyr: 20 mM pyruvate; Fru: 20 mM fructose; Suc: 15 mM succinate; Lac: 20 mM lactose; Ara: 20 mM arabinose; Mal: 20 mM maltose; and Glc: 0.4% v/v glucose. For all minimal media experiments, expression levels were extracted between OD 0.2 and 0.8, during which growth rate remained approximately constant. Protein concentration is also measured at the single-cell level via fluorescence microscopy (filled points). Filled points represent the means of the Tar concentration distributions shown in EDF 4. Hollow points are the means of 8 technical replicates for each carbon source in minimal media or 40 technical replicates for each OD in rich media. Solid lines are fits to the data. Growth rate, OD, and batch fluorescence levels were assessed in the same batch cultures using a plate reader. (Bottom) Same as the top panel, but for Tsr concentration. Filled points represent the means of the Tsr concentration distributions shown in EDF 4. Inset: growth rate as a function of OD. Points represent growth rates extracted at the same culture phases as expression levels (see methods). **(b)** Scaling of mean Tar/Tsr receptor ratio as a function of growth rate for cells grown in rich (circles) and minimal (squares) media. Same color coding as figure 2c and e. Error bars indicate 95% confidence intervals of the mean *K*_1*/*2_ obtained via bootstrap resampling and are typically on the order of the data point size. **(c)** Scaling of mean L-serine (black) and L-aspartate (blue) *K*_1*/*2_ as a function of growth rate for cells grown in rich and minimal media. Error bars indicate 95% confidence intervals of the mean *K*_1*/*2_ obtained via bootstrap resampling and are typically on the order of the data point size. **(d)** Coefficient of variation (CV) of L-serine (black) and L-aspartate (blue) *K*_1*/*2_ as a function of growth rate. Error bars indicate 95% confidence intervals of the CV obtained via bootstrap resampling. **(e)** CV of tumble bias (green) and run speed (brown) as a function of growth rate. Error bars indicate 95% confidence intervals of the CV obtained via bootstrap resampling and are typically on the order of the data point size.

Taken together, these results suggest that growth rate is the primary — if not sole — determinant of the chemoreceptor ratio. Differences in nutrient availability across growth conditions (due to nutrient consumption at varying cell densities in rich media or different carbon sources in minimal media) modulate growth rate, which in turn regulates chemoreceptor concentrations and thus the Tar/Tsr receptor ratio (Figure 5b). This regulation ultimately translates non-linearly into ligand sensitivities (Figure 5c and EDF 8). The non-linear mapping between receptor ratio and *K*_1*/*2_ when switching from rich to minimal media likely arises from differences in receptor cluster organization rather than cluster protein stoichiometry. Cryo-electron tomography data have shown that cells grown in minimal media form more tightly-packed receptor clusters compared to cells grown in rich media, despite the similar overall cluster sizes [55]. However, the receptor-to-CheA and receptor-to-CheW stoichiometry is conserved across these two conditions [33, 55], likely because the mocha operon (which includes *cheW* and *cheA*) and meche operon (which includes *tar*), as well as the *tsr* operon are all regulated by FliA [25].

We also found that the degree of sensory diversity, characterized by the CV of *K*_1*/*2_, varies as a function of growth rate (Figure 5d). Interestingly, the trends for aspartate and serine are opposite in cells grown in rich media: at fast growth rates, cells exhibit reduced diversity in their sensitivity to serine and increased diversity in their sensitivity to aspartate, with this pattern reversing at slow growth rates. These trends are consistent with a bet-hedging strategy in which bacterial populations suppress diversity in sensing current resources for focused exploitation, while increasing diversity in sensing gradients not yet encountered to boost exploration. Indeed, quantification of L-serine and L-aspartate concentrations in rich media as a function of cell density revealed that *E. coli* preferentially consumes serine at low cell densities (corresponding to fast growth) before switching to aspartate consumption at higher cell densities (slow growth) (EDF 9a) [39–41]—meaning that cells narrow their chemotactic diversity for the exact ligand being actively consumed, while broadening it for the one not yet exploited. In line with this strategy, *K*_1*/*2_ reaches its lowest value in rich media (corresponding to the highest sensitivity) when cells are actively consuming the corresponding ligand (Figure 1g), suggesting an anticipatory mechanism that primes cells to detect alternative nutrient sources once the current one is depleted. In environments of greater uncertainty—such as minimal media, where we found that neither serine nor aspartate is present (EDF 9b) and the growth rate remains constant during exponential growth—cells maintain uniformly high chemotactic diversity for both amino acids, ensuring readiness to detect either when it becomes available.

Finally, we wondered whether there is a similar bet-hedging mechanism at the level of motile behavior. We quantified the two key parameters that define the random walk of bacteria in absence of chemical gradients: tumble bias and run speed. These parameters have been shown to vary greatly within isogenic populations of *E. coli* cells [28]. We tracked the swimming behavior of motile, wild-type cells—the parent strain used for our FRET-based sensory diversity measurements—using a microfluidic device that allows tracking hundreds of cells simultaneously in a quasi-two-dimensional isotropic environment using phase-contrast microscopy [34]. As expected, we observed substantial diversity in run speed (CV ∼ 0.25) and even greater variability in tumble bias (CV∼ 1). However, neither the diversity in run speed nor in tumble bias changed as a function of growth rate (Figure 5e). Similarly, the mean values of both run speed and tumble bias showed only weak dependence on growth rate (EDF 10). The latter observation is consistent with our recent findings in a different *E. coli* strain, RP437, where we reported a constant mean tumble bias across a range of optical densities [34], and it also agrees with an earlier study that reported that the swimming speed of strain HE206 varies only marginally with growth rate [56]. However, it contrasts with older reports describing a non-monotonic relationship between mean run speed and optical density [57]. This discrepancy likely arises from differences in the strains used: RP437 in this earlier study, versus MG1655 here and HE206 in [56].

### 2.7 Chemosensory diversity boosts collective navigation in contrasting environments via phenotypic filtering

Does the increased sensory diversity we observe under some environments in Figure 5d actually confer a behavioral advantage? Or is the mean sensory phenotype the only important parameter for performance? To address this question, we devised a bacterial wave speed competition experiment of two populations with the same mean receptor ratio but different receptor ratio variances. To this end, we utilized motile versions of the wild-type and high-variability populations described in Figure 3, each expressing cytosolic YFP to facilitate wave tracking. To decouple sensing from growth, we suspended the cells in a medium that arrests growth, effectively keeping their receptor ratio constant throughout the experiment (see methods). We introduced the two populations in separate microfluidic swim channels [58] containing either 100 *µ*M L-aspartate or 100 *µ*M L-serine (Figure 6a). The cells rapidly formed chemotactic waves by creating a gradient of ligand through consumption and directed movement (Figure 6b) [21, 58, 59]. Both populations migrated faster in serine than aspartate, as it has been shown before [22, 34, 60]. Comparison of the wave velocities of wild-type and high-variability populations revealed a substantial advantage of the high-variability population in both aspartate and serine gradients (Figures 6c and EDF 6). This advantage was not due to differences in individual motility traits, as both populations exhibited identical tumble bias and run speed distributions (EDF 6). These findings demonstrate a clear evolutionary advantage of increased sensory diversity, potentially enabling more diverse populations to colonize new territories faster, thus improving their overall chance of survival in different environments.

**Fig. 6.**
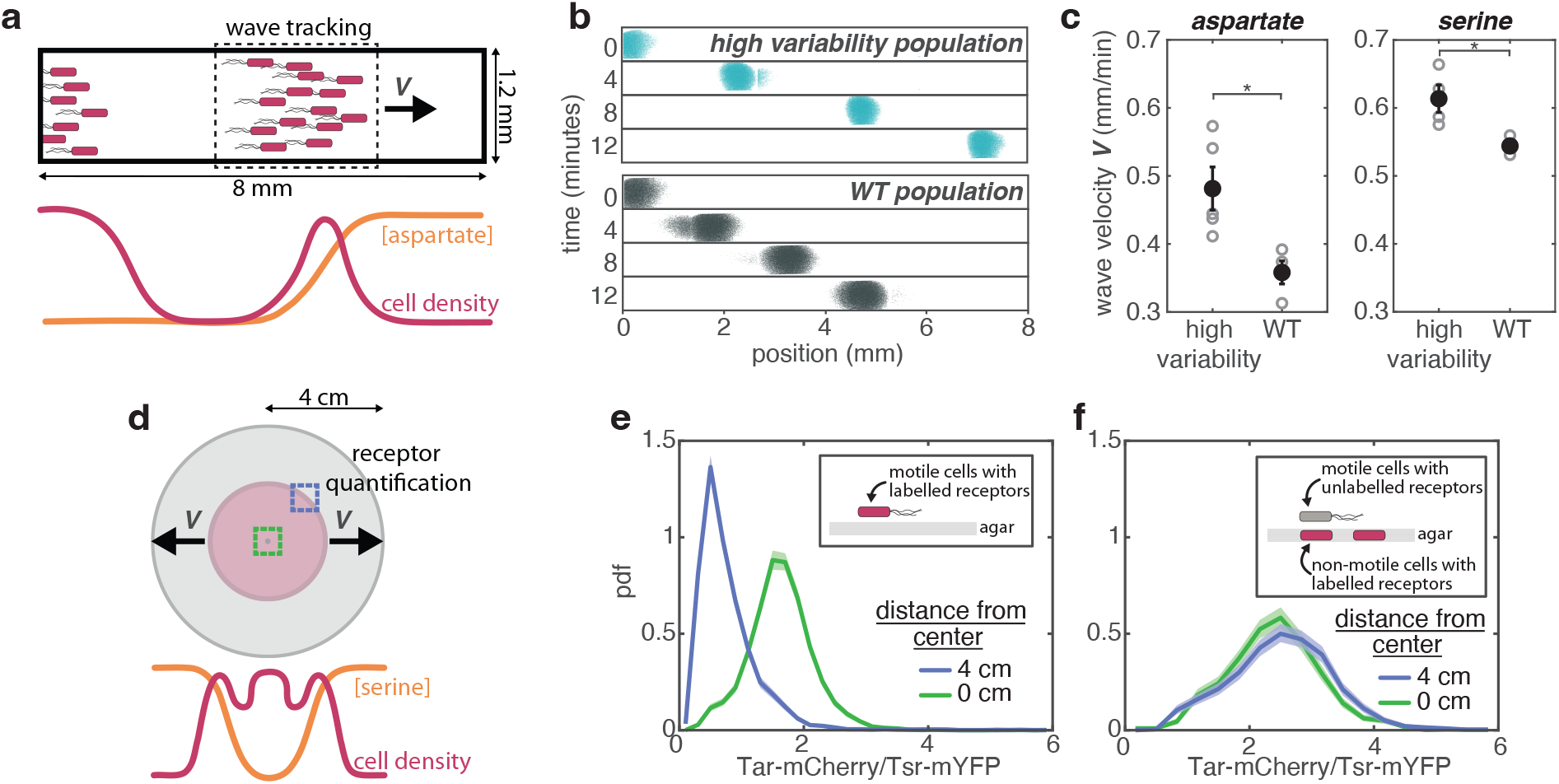
Sensory diversity enhances the chemotactic migration of isogenic populations across environments through phenotypic filtering **(a)** Schematic of the bacterial speed competition assay. Cells introduced at the entrance of a microfluidic channel form a band that migrates through a self-generated gradient of L-aspartate, established by bacterial consumption. Segmented images of bacterial bands formed by a high variability population (top) and a wild-type (WT) population (bottom). Images are from two separate experiments, aligned at time t = 0 for comparison. Refer to Figures 3b and 3c for the Tar/Tsr ratio and *K*_1*/*2_ distributions of these two populations, respectively. **(c)** Wave velocities of cells migrating up a 100*µ*M L-aspartate gradient (left) or a 100*µ*M L-serine gradient (right). Hollow points represent independent biological replicates, while solid points indicate the mean values across all replicates. Error bars represent standard error of the mean. Means are statistically significant different, as determined by a two-sample t-test with p = 0.0159 for aspartate waves and p = 0.0385 for the serine waves. **(d)** Schematic of the swim plate assay. Cells on rich media swim plates form a high-density band that migrates up a self-generated L-serine gradient. **(e)** Histograms of the Tar/Tsr chemoreceptor ratio, measured via fluorescence microscopy for motile cells collected from the edge (blue) or center (green) of the expanding colony. **(f)** Same as panel e, but for non-motile cells acting as a biosensor for gene expression collected at the edge (blue) or center (green) of the colony. A motile but unlabeled strain was used to establish the chemical gradient experienced by the biosensor cells. Solid lines represent the histograms of all Tar/Tsr ratios from a single experiment. Shaded areas indicate 95% confidence intervals obtained through bootstrap resampling.

To explore the exact mechanism by which high-variability populations climb a ligand gradient faster, we employed the classical chemotaxis assay on soft agar swim plates [61] using motile cells with labeled Tar and Tsr receptors (Figure 6d). On a soft agar swim plate containing nutrient-rich media, *E. coli* cells form a colony with multiple bands of high cell density, the first of which consumes serine [61]. We collected cells from the serine-consuming band and measured their receptor ratios using fluorescence microscopy. At the same time, we measured the receptor ratios of cells in the center of the colony, near the inoculation point. We observed a striking difference between these two populations: cells from the serine-consuming band had a receptor ratio biased towards higher Tsr and lower Tar, compared to those from the center of the colony (Figure 6e), demonstrating the existence of a receptor ratio sorting mechanism. This finding aligns with our previous theoretical prediction of such sorting mechanism [38] and its experimental demonstration on Tsr selection on swim plates containing a single amino acid [34]. This sorting mechanism is non-genetic, as demonstrated in an experiment using a non-motile strain with labeled Tar and Tsr receptors acting as a biosensor for gene expression. We spread the biosensor strain uniformly across the agar matrix and used its unlabeled wild-type parent strain to establish chemoattractant gradients on the plate. By measuring the receptor ratio of the non-motile strain, we confirmed that there are no changes in the expression of the Tar and Tsr receptors at the center and edge of the colony (Figure 6f), consistent with our previous findings for Tsr expression [34].

During collective migration, this non-genetic sorting mechanism likely enriches the migrating band with cells best suited for climbing attractant gradients: cells with a low Tar/Tsr ratio are selected for in serine gradients, while cells with a high Tar/Tsr ratio are selected for in aspartate gradients. The high-variability population contains cells with both higher and lower Tar/Tsr ratios than the wild-type population. As a result, the sorting mechanism can adapt the phenotypic composition of the migrating band further in the case of the high-variability population, resulting in better chemotactic performance. Since the sorting mechanism is non-genetic, it enables rapid phenotypic filtering even in the absence of growth, as it is the case in our bacterial wave speed competition experiment.

## 3 Discussion

Phenotypic heterogeneity in isogenic bacterial populations was first noted more than 80 years ago in the context of antibiotic persistence [62, 63]. Within the bacterial chemotaxis system studied here, Spudich and Koshland demonstrated already 50 years ago that isogenic cell populations exhibit substantial behavioral diversity [24]. The advent of single-cell technologies in the last two decades have enabled systematic studies of these phenomena as bet-hedging strategies [6, 27, 31, 64–66], but most have focused on benefits for growth and survival at the single-cell level. In this study, we systematically investigated the mechanisms and consequences of phenotypic diversity in *E. coli* chemotaxis across scales, from stochastic gene expression to population-level collective navigation, under various environmental conditions.

At the intracellular level, we pinpointed the origin of sensory diversity to a single cellular random variable: the ratio between the stochastic expression levels of the two major chemoreceptors Tar and Tsr. We further demonstrated that diversity in the Tar/Tsr ratio, and hence also the sensitivity to chemoeffector ligands, depends on the growth environment. By contrast, we found that diversity in the run-and-tumble motile behavior itself remains constant across environments. The environment-dependent modulation of sensory diversity appears to stem largely from the dependence of *tar* expression on growth rate, whereas *tsr* expression remains nearly constant. Consequently, cells modulate not only the mean ligand sensitivity but also its variance in a manner that is tied to growth rate, without requiring dedicated regulatory pathways that couple sensing of specific nutrients to gene expression. Given that *tar* and *tsr* are located on separate operons, it seems likely that this differential regulation occurs at the level of transcription, although our present experiments do not rule out the possibility of post-transcriptional mechanisms.

Operon structure also likely explains the invariance of tumble bias distributions across different growth conditions. Tumble bias is controlled by the expression ratio of two pairs of chemotaxis proteins CheY-CheZ and CheR-CheB [25, 67, 68]. The genes encoding these four proteins are located on the same operon as *tar* (the meche operon), transcribing a single polycistronic mRNA molecule. This organization enforces strong transcriptional [25] and translational [68] coupling between these four proteins and ensures that while the absolute copy numbers of these proteins may vary with growth rate — similar to how Tar copy numbers change (Figure 5a) — their relative ratios remain conserved [25, 68]. On the other hand, the primary determinants of run speed are flagellar number and flagellar length [69]. These parameters appear to remain constant over the growth rate range examined here [56]. This is likely due to the growthrate independent expression of FliA (*σ*^28^), the sigma factor responsible for regulating the expression of flagellar filament and motor genes [56]. Thus, while the diversity of sensory response is modulated by the growth environment, we found that key parameters of the motility machinery remained invariant.

Beyond shaping sensory diversity, environmental regulation of the Tar/Tsr ratio is also expected to control temporal noise strength. As we have recently shown [70], low Tar/Tsr ratios increase noise relative to intermediate ratios by shifting the receptor cluster closer to a critical point.

Our tests of the consequence of sensory diversity yielded the surprising finding that, beyond single-cell chemotaxis, it can implement bet-hedging at the level of collective migration phenotypes. The hypothesis that *E. coli* exploits sensory diversity to enhance chemotactic navigation [31] has been tested at the single-cell level in agent-based simulations [8], and are consistent with earlier experimental results on single-cell gradient responses in a microfluidic maze [30]. But collective chemotactic migrations require that all cells within the advancing traveling wave of population density to share the same chemotactic drift velocity [21] and hence requires substantial population-level coordination. Naively, a high degree of sensory diversity would seem detrimental to such coordination, given that ligand response sensitivity is a key parameter controlling chemotactic drift velocity [21, 71]. We resolved this apparent paradox by examining gene expression within the migrating collectives. Remarkably, those experiments revealed how cells with a gene expression pattern favorable to fast chemotactic migration (specifically, a low Tar/Tsr ratio within the tested environment with serine as the chemoeffector) are selected by the collective migration behavior itself. The latter phenomenon of phenotype selection within collectively migrating populations was recently predicted [38], and experimentally verified [34] as a mechanism for non-genetic adaptation of population diversity - effectively modulating both the mean and variance of phenotype distributions.

Our results in the present study demonstrate that when combined with environmentally modulated stochastic gene expression, non-genetic diversity adaptation can enable bet-hedging strategies not only at the level of single-cell phenotypes but also for population-level collective behaviors. Moreover, we demonstrated that the environmental modulation of sensory diversity operates through growth rate as a control variable. This enables a collective chemotactic bet-hedging strategy that suppresses response diversity toward current growth resources for efficient exploitation, while boosting it toward as yet unencountered resources to enhance exploration. This bet-hedging strategy ensures that in rapidly fluctuating environments, such as terrestrial soils [72] or the mammalian gut [73], a subset of the population is always pre-adapted, priming the entire population to detect and collectively exploit emerging nutrient sources as they become available.

## 4 Acknowledgments

We thank Simone Boskamp for her invaluable assistance with cloning and transformations and Dimitry Lamers for assisting with the microfluidic chemostat wafer fabrication. We also thank Lam Vo for providing the microfluidic device for wave tracking, Jyot Antani for helping set up the cell tracking experiment, and Gustavo Madeira Santana for helping set up the FRET experiment. We thank Marko Kamp for microscopy support and Brahim Ait Said for software support. We thank Victor Sourjik and Howard Berg for providing plasmids and strains that were used in this study. Finally, we thank John S. Parkinson, Jonas Cremer, Keita Kamino, Jeremy Moore, Lam Vo, Diana Valverde Mendez, Evan Usher, and Johannes Keegstra for helpful discussions.

## 5 Author contributions

FA, TSS, and TE conceived the study. FA performed experiments. RJ assisted with receptor quantification experiments. FA analyzed the data and wrote the manuscript. FA, RJ, TSS, and TE edited the manuscript. This work was supported by NIGMS awards R01GM106189 (TE and TSS), R01GM138533 and R35GM158058 (TE), and by the Alfred P. Sloan Foundation G-2023-19668 (FA and TE).

## Notes

### Competing Interest Statement

The authors have declared no competing interest.

### Summary of Updates

The manuscript has been updated to include new data (measurements of amino acid concentrations). The funding information has also been updated.

## References

[1] Keegstra, J.M., Carrara, F., Stocker, R.: The ecological roles of bacterial chemotaxis. Nature Reviews Microbiology 20(8), 491–504 (2022) 10.1038/s41579-022-00709-w

[2] Ackermann, M.: A functional perspective on phenotypic heterogeneity in microorganisms. Nature Reviews Microbiology 13(8), 497–508 (2015) 10.1038/nrmicro3491

[3] Xue, B., Sartori, P., Leibler, S.: Environment-to-phenotype mapping and adaptation strategies in varying environments. Proceedings of the National Academy of Sciences 116(28), 13847–13855 (2019) 10.1073/pnas.1903232116

[4] Carey, J.N., Mettert, E.L., Roggiani, M., Myers, K.S., Kiley, P.J., Goulian, M.: Regulated Stochasticity in a Bacterial Signaling Network Permits Tolerance to a Rapid Environmental Change. Cell 173(1), 196–20714 (2018) 10.1016/j.cell.2018.02.005

[5] Schreiber, F., Littmann, S., Lavik, G., Escrig, S., Meibom, A., Kuypers, M.M.M., Ackermann, M.: Phenotypic heterogeneity driven by nutrient limitation promotes growth in fluctuating environments. Nature Microbiology 1(6), 16055 (2016) 10.1038/nmicrobiol.2016.55

[6] Frankel, N.W., Pontius, W., Dufour, Y.S., Long, J., Hernandez-Nunez, L., Emonet, T.: Adaptability of non-genetic diversity in bacterial chemotaxis. eLife 3, 03526 (2014) 10.7554/eLife.03526

[7] Waite, A.J., Frankel, N.W., Emonet, T.: Behavioral Variability and Phenotypic Diversity in Bacterial Chemotaxis. Annual Review of Biophysics 47(1), 595–616 (2018) 10.1146/annurev-biophys-062215-010954

[8] Moore, J.P., Kamino, K., Kottou, R., Shimizu, T.S., Emonet, T.: Signal integration and adaptive sensory diversity tuning in Escherichia coli chemotaxis. Cell Systems 15(7), 628–6388 (2024) 10.1016/j.cels.2024.06.003

[9] Miller, M.B., Bassler, B.L.: Quorum sensing in bacteria. Annual Review of Microbiology 55(1), 165–199 (2001) 10.1146/annurev.micro.55.1.165

[10] Pai, A., You, L.: Optimal tuning of bacterial sensing potential. Molecular Systems Biology 5(1), 286 (2009) 10.1038/msb.2009.43

[11] Flemming, H.-C., Wingender, J.: The biofilm matrix. Nature Reviews Microbiology 8(9), 623–633 (2010) 10.1038/nrmicro2415

[12] Nadell, C.D., Foster, K.R., Xavier, J.B.: Emergence of spatial structure in cell groups and the evolution of cooperation. PLoS Computational Biology 6(3), 1000716 (2010) 10.1371/journal.pcbi.1000716

[13] Lee, H.H., Molla, M.N., Cantor, C.R., Collins, J.J.: Bacterial charity work leads to population-wide resistance. Nature 467(7311), 82–85 (2010) 10.1038/nature09354

[14] Yurtsev, E.A., Chao, H.X., Datta, M.S., Artemova, T., Gore, J.: Bacterial cheating drives the population dynamics of cooperative antibiotic resistance plasmids. Molecular Systems Biology 9(1), 683 (2013) 10.1038/msb.2013.39

[15] Srinivasan, S., Vladescu, I.D., Koehler, S.A., Wang, X., Mani, M., Rubinstein, S.M.: Matrix production and sporulation in bacillus subtilis biofilms localize to propagating wave fronts. Biophysical Journal 114(6), 1490–1498 (2018) 10.1016/j.bpj.2018.02.002

[16] Chou, K.-T., Lee, D.-Y.D., Chiou, J.-G., Galera-Laporta, L., Ly, S., Garcia-Ojalvo, J., Süel, G.M.: A segmentation clock patterns cellular differentiation in a bacterial biofilm. Cell 185(1), 145–157 (2022) 10.1016/j.cell.2021.12.001

[17] Kaiser, D.: Coupling cell movement to multicellular development in myxobacteria. Nature Reviews Microbiology 1(1), 45–54 (2003) 10.1038/nrmicro733

[18] Saulnier, J.-B., Romanos, M., Schrohe, J., Cuzin, C., Calvez, V., Mignot, T.: The mechanism of spatial pattern transition in motile bacterial collectives. bioRxiv (2024) 10.1101/2024.10.28.620572

[19] Rutherford, S.T., Bassler, B.L.: Bacterial quorum sensing: Its role in virulence and possibilities for its control. Cold Spring Harbor Perspectives in Medicine 2(11), 012427 (2012) 10.1101/cshperspect.a012427

[20] Cornforth, D.M., Foster, K.R.: Competition sensing: The social side of bacterial stress responses. Nature Reviews Microbiology 11(4), 285–293 (2013) 10.1038/nrmicro2977

[21] Fu, X., Kato, S., Long, J., Mattingly, H.H., He, C., Vural, D.C., Zucker, S.W., Emonet, T.: Spatial self-organization resolves conflicts between individuality and collective migration. Nature Communications 9(1), 2177 (2018) 10.1038/s41467-018-04539-4

[22] Cremer, J., Honda, T., Tang, Y., Wong-Ng, J., Vergassola, M., Hwa, T.: Chemotaxis as a navigation strategy to boost range expansion. Nature 575(7784), 658–663 (2019) 10.1038/s41586-019-1733-y

[23] Gude, S., Pinçe, E., Taute, K.M., Seinen, A.-B., Shimizu, T.S., Tans, S.J.: Bacterial coexistence driven by motility and spatial competition. Nature 578(7796), 588–592 (2020) 10.1038/s41586-020-2033-2

[24] Spudich, J.L., Koshland, D.E.: Non-genetic individuality: chance in the single cell. Nature 262(5568), 467–471 (1976) 10.1038/262467a0

[25] Kollmann, M., Løvdok, L., Bartholomé, K., Timmer, J., Sourjik, V.: Design principles of a bacterial signalling network. Nature 438(7067), 504–507 (2005) 10.1038/nature04228

[26] Park, H., Pontius, W., Guet, C.C., Marko, J.F., Emonet, T., Cluzel, P.: Interdependence of behavioural variability and response to small stimuli in bacteria. Nature 468(7325), 819–823 (2010) 10.1038/nature09551

[27] Keegstra, J.M., Kamino, K., Anquez, F., Lazova, M.D., Emonet, T., Shimizu, T.S.: Phenotypic diversity and temporal variability in a bacterial signaling network revealed by single-cell FRET. eLife 6, 27455 (2017) 10.7554/eLife.27455

[28] Waite, A.J., Frankel, N.W., Dufour, Y.S., Johnston, J.F., Long, J., Emonet, T.: Non-genetic diversity modulates population performance. Molecular Systems Biology 12(12), 895 (2016) 10.15252/msb.20167044

[29] Dufour, Y.S., Fu, X., Hernandez-Nunez, L., Emonet, T.: Limits of Feedback Control in Bacterial Chemotaxis. PLoS Computational Biology 10(6), 1003694 (2014) 10.1371/journal.pcbi.1003694

[30] Salek, M.M., Carrara, F., Fernandez, V., Guasto, J.S., Stocker, R.: Bacterial chemotaxis in a microfluidic T-maze reveals strong phenotypic heterogeneity in chemotactic sensitivity. Nature Communications 10(1), 1877 (2019) 10.1038/s41467-019-09521-2

[31] Kamino, K., Keegstra, J.M., Long, J., Emonet, T., Shimizu, T.S.: Adaptive tuning of cell sensory diversity without changes in gene expression. Science Advances 6(46), 1087 (2020) 10.1126/sciadv.abc1087

[32] Parkinson, J.S., Hazelbauer, G.L., Falke, J.J.: Signaling and sensory adaptation in Escherichia coli chemoreceptors: 2015 update. Trends in Microbiology 23(5), 257–266 (2015) 10.1016/j.tim.2015.03.003

[33] Li, M., Hazelbauer, G.L.: Cellular Stoichiometry of the Components of the Chemotaxis Signaling Complex. Journal of Bacteriology 186(12), 3687–3694 (2004) 10.1128/JB.186.12.3687-3694.2004

[34] Vo, L., Avgidis, F., Mattingly, H.H., Edmonds, K., Burger, I., Balasubramanian, R., Shimizu, T.S., Kazmierczak, B.I., Emonet, T.: Nongenetic adaptation by collective migration. Proceedings of the National Academy of Sciences 122(8), 2423774122 (2025) 10.1073/pnas.2423774122

[35] Monod, J., Wyman, J., Changeux, J.-P.: On the nature of allosteric transitions: A plausible model. Journal of Molecular Biology 12(1), 88–118 (1965) 10.1016/S0022-2836(65)80285-6

[36] Mello, B.A., Tu, Y.: An allosteric model for heterogeneous receptor complexes: Understanding bacterial chemotaxis responses to multiple stimuli. Proceedings of the National Academy of Sciences 102(48), 17354–17359 (2005) 10.1073/pnas.0506961102

[37] Wang, P., Robert, L., Pelletier, J., Dang, W.L., Taddei, F., Wright, A., Jun, S.: Robust Growth of Escherichia coli. Current Biology 20(12), 1099–1103 (2010) 10.1016/j.cub.2010.04.045

[38] Mattingly, H.H., Emonet, T.: Collective behavior and nongenetic inheritance allow bacterial populations to adapt to changing environments. Proceedings of the National Academy of Sciences 119(26), 2117377119 (2022) 10.1073/pnas.2117377119

[39] Yang, Y., M. Pollard, A., Höfler, C., Poschet, G., Wirtz, M., Hell, R., Sourjik, V.: Relation between chemotaxis and consumption of amino acids in bacteria. Molecular Microbiology 96(6), 1272–1282 (2015) 10.1111/mmi.13006

[40] Prüss, B.M., Nelms, J.M., Park, C., Wolfe, A.J.: Mutations in NADH:ubiquinone oxidoreductase of Escherichia coli affect growth on mixed amino acids. Journal of Bacteriology 176(8), 2143–2150 (1994) 10.1128/jb.176.8.2143-2150.1994

[41] Selvarasu, S., Ow, D.S., Lee, S.Y., Lee, M.M., Oh, S.K., Karimi, I.A., Lee, D.: Characterizing Escherichia coli DH5 growth and metabolism in a complex medium using genome-scale flux analysis. Biotechnology and Bioengineering 102(3), 923–934 (2009) 10.1002/bit.22119

[42] Adler, J., Hazelbauer, G.L., Dahl, M.M.: Chemotaxis Toward Sugars in Escherichia coli. Journal of Bacteriology 115(3), 824–847 (1973) 10.1128/jb.115.3.824-847.1973

[43] Taniguchi, Y., Choi, P.J., Li, G.-W., Chen, H., Babu, M., Hearn, J., Emili, A., Xie, X.S.: Quantifying E. coli Proteome and Transcriptome with Single-Molecule Sensitivity in Single Cells. Science 329(5991), 533–538 (2010) 10.1126/science.1188308

[44] Kalinin, Y., Neumann, S., Sourjik, V., Wu, M.: Responses of Escherichia coli Bacteria to Two Opposing Chemoattractant Gradients Depend on the Chemoreceptor Ratio. Journal of Bacteriology 192(7), 1796–1800 (2010) 10.1128/JB.01507-09

[45] Li, L., Zhang, X., Sun, Y., Ouyang, Q., Tu, Y., Luo, C.: Phenotypic Variability Shapes Bacterial Responses to Opposing Gradients. PRX Life 2(1), 013001 (2024) 10.1103/PRXLife.2.013001

[46] Frank, V., Livne, N., Koler, M., Vaknin, A.: Single-array measurements reveal non-uniform, mosaic-like chemosensory arrays in bacteria. Nature Communications 17(1), 587 (2026) 10.1038/s41467-025-67285-4

[47] Yoney, A., Salman, H.: Precision and Variability in Bacterial Temperature Sensing. Biophysical Journal 108(10), 2427–2436 (2015) 10.1016/j.bpj.2015.04.016

[48] Salman, H., Libchaber, A.: A concentration-dependent switch in the bacterial response to temperature. Nature Cell Biology 9(9), 1098–1100 (2007) 10.1038/ncb1632

[49] Yang, Y., Sourjik, V.: Opposite responses by different chemoreceptors set a tunable preference point in escherichia coli ph taxis. Molecular Microbiology 86(6), 1482–1489 (2012) 10.1111/mmi.12070

[50] Chilcott, G.S., Hughes, K.T.: Coupling of Flagellar Gene Expression to Flagellar Assembly in Salmonella enterica Serovar Typhimurium and Escherichia coli. Microbiology and Molecular Biology Reviews 64(4), 694–708 (2000) 10.1128/MMBR.64.4.694-708.2000

[51] Elowitz, M.B., Levine, A.J., Siggia, E.D., Swain, P.S.: Stochastic Gene Expression in a Single Cell. Science 297(5584), 1183–1186 (2002) 10.1126/science.1070919

[52] Balakrishnan, R., Mori, M., Segota, I., Zhang, Z., Aebersold, R., Ludwig, C., Hwa, T.: Principles of gene regulation quantitatively connect DNA to RNA and proteins in bacteria. Science 378(6624), 2066 (2022) 10.1126/science.abk2066

[53] Ames, P., Studdert, C.A., Reiser, R.H., Parkinson, J.S.: Collaborative signaling by mixed chemoreceptor teams in Escherichia coli. Proceedings of the National Academy of Sciences 99(10), 7060–7065 (2002) 10.1073/pnas.092071899

[54] Ni, B., Colin, R., Link, H., Endres, R.G., Sourjik, V.: Growth-rate dependent resource investment in bacterial motile behavior quantitatively follows potential benefit of chemotaxis. Proceedings of the National Academy of Sciences 117(1), 595–601 (2020) 10.1073/pnas.1910849117

[55] Khursigara, C.M., Lan, G., Neumann, S., Wu, X., Ravindran, S., Borgnia, M.J., Sourjik, V., Milne, J., Tu, Y., Subramaniam, S.: Lateral density of receptor arrays in the membrane plane influences sensitivity of the E. coli chemotaxis response. The EMBO Journal 30(9), 1719–1729 (2011) 10.1038/emboj.2011.77

[56] Honda, T., Cremer, J., Mancini, L., Zhang, Z., Pilizota, T., Hwa, T.: Coordination of gene expression with cell size enables Escherichia coli to efficiently maintain motility across conditions. Proceedings of the National Academy of Sciences 119(37), 2110342119 (2022) 10.1073/pnas.2110342119

[57] Staropoli, J.F., Alon, U.: Computerized Analysis of Chemotaxis at Different Stages of Bacterial Growth. Biophysical Journal 78(1), 513–519 (2000) 10.1016/S0006-3495(00)76613-6

[58] Phan, T.V., Mattingly, H.H., Vo, L., Marvin, J.S., Looger, L.L., Emonet, T.: Direct measurement of dynamic attractant gradients reveals breakdown of the Patlak–Keller–Segel chemotaxis model. Proceedings of the National Academy of Sciences 121(3), 2309251121 (2024) 10.1073/pnas.2309251121

[59] Bai, Y., He, C., Chu, P., Long, J., Li, X., Fu, X.: Spatial modulation of individual behaviors enables an ordered structure of diverse phenotypes during bacterial group migration. eLife 10, 67316 (2021) 10.7554/eLife.67316

[60] Wong-Ng, J., Melbinger, A., Celani, A., Vergassola, M.: The Role of Adaptation in Bacterial Speed Races. PLOS Computational Biology 12(6), 1004974 (2016) 10.1371/journal.pcbi.1004974

[61] Wolfe, A.J., Berg, H.C.: Migration of bacteria in semisolid agar. Proceedings of the National Academy of Sciences 86(18), 6973–6977 (1989) 10.1073/pnas.86.18.6973

[62] Hobby, G.L., Meyer, K., Chaffee, E.: Observations on the mechanism of action of penicillin. Proceedings of the Society for Experimental Biology and Medicine 50(2), 281–285 (1942) 10.3181/00379727-50-13773

[63] Bigger, J.W.: Treatment of staphylococcal infections with penicillin by intermittent sterilisation. The Lancet 244(6320), 497–500 (1944) 10.1016/S0140-6736(00)74210-3

[64] Balaban, N.Q., Merrin, J., Chait, R., Kowalik, L., Leibler, S.: Bacterial Persistence as a Phenotypic Switch. Science 305(5690), 1622–1625 (2004) 10.1126/science.1099390

[65] Kussell, E., Leibler, S.: Phenotypic Diversity, Population Growth, and Information in Fluctuating Environments. Science 309(5743), 2075–2078 (2005) 10.1126/science.1114383

[66] Veening, J.-W., Stewart, E.J., Berngruber, T.W., Taddei, F., Kuipers, O.P., Hamoen, L.W.: Bet-hedging and epigenetic inheritance in bacterial cell development. Proceedings of the National Academy of Sciences 105(11), 4393–4398 (2008) 10.1073/pnas.0700463105

[67] Dufour, Y.S., Gillet, S., Frankel, N.W., Weibel, D.B., Emonet, T.: Direct Correlation between Motile Behavior and Protein Abundance in Single Cells. PLOS Computational Biology 12(9), 1005041 (2016) 10.1371/journal.pcbi.1005041

[68] Løvdok, L., Bentele, K., Vladimirov, N., Müller, A., Pop, F.S., Lebiedz, D., Kollmann, M., Sourjik, V.: Role of Translational Coupling in Robustness of Bacterial Chemotaxis Pathway. PLoS Biology 7(8), 1000171 (2009) 10.1371/journal.pbio.1000171

[69] Lisevich, I., Colin, R., Yang, H.Y., Ni, B., Sourjik, V.: Physics of swimming and its fitness cost determine strategies of bacterial investment in flagellar motility. Nature Communications 16(1), 1731 (2025) 10.1038/s41467-025-56980-x

[70] Keegstra, J.M., Avgidis, F., Usher, E., Mulla, Y., Parkinson, J.S., Shimizu, T.S.: Spontaneous switching in a protein signalling array reveals near-critical cooperativity. Nature Physics (2026) 10.1038/s41567-025-03158-3. Published 29 January 2026

[71] Si, G., Wu, T., Ouyang, Q., Tu, Y.: Pathway-based mean-field model for escherichia coli chemotaxis. Physical Review Letters 109(4), 048101 (2012) 10.1103/PhysRevLett.109.048101

[72] Young, I.M., Crawford, J.W.: Interactions and self-organization in the soil-microbe complex. Science 304(5677), 1634–1637 (2004) 10.1126/science.1097394

[73] Donaldson, G.P., Lee, S.M., Mazmanian, S.K.: Gut biogeography of the bacterial microbiota. Nature Reviews Microbiology 14(1), 20–32 (2016) 10.1038/nrmicro3552

