## Supplementary Information for "Growth-dependent sensory bet-hedging enhances collective navigation"

#### 1 Strains and plasmids

*E. coli* strain MG1655 ΔFliC::FLP, a generous gift from Victor Sourjik, was used as a background for all FRET and receptor quantification experiments. For FRET experiments, we conducted an in-frame deletion of the CheRBYZ genes. Subsequently, the gene expressing a glass-adhesive mutant of FliC [1] was cloned at the native FliC locus using homologous recombination, resulting in strain TSS2191. The FRET acceptor-donor pair, consisting of CheY-mRFP1 and CheZ-mYFP (A206K variant), was expressed in tandem from a pTrc99A plasmid under IPTG induction (pSJAB106, [1]). To determine the optimal levels of IPTG induction, we compared the motility of strain TSS2114/pSJAB106 to that of WT strain TSS2096 carrying an empty FRET plasmid (pTrc99A) on soft agar swim plates (EDF 1).

For receptor quantification experiments, *tsr* was translationally fused with mYFP (A206K variant), and *tar* was translationally fused with mCherry, at their respective native chromosomal loci using the same MG1655 ΔFliC::FLP parent strain as in FRET experiments. The cheRB genes were deleted in-frame (yielding strain TSS2155), ensuring comparable Tar receptor expression to that of the strain used for FRET (TSS2191). Since the cheR and cheB genes are located in the same operon as the *tar* gene, factors such as the primary transcript mRNA length can affect the protein copy numbers of Tar.

For FRET experiments involving receptor overexpression, we constructed a high variability strain by transforming TSS2191 with a pTrc99A plasmid expressing wild-type Tar under IPTG induction (pSJAB21). The same FRET acceptor-donor pair used in all other FRET experiments was cloned into a pRZ30 plasmid under NaSal induction (pSJAB196). To quantify receptor overexpression, strain TSS2155 was transformed with a pTrc99A plasmid expressing a Tar-mCherry fusion under IPTG induction (pMV13). Since pSJAB21 and pMV13 are based on the same plasmid backbone, pTrc99A, and are induced at the same level, wild-type Tar and Tar-mCherry are expected to exhibit uniform expression levels.

For swimming competition experiments, wild-type MG1655, a generous gift from Victor Sourjik, was transformed with the same IPTG-inducible pTrc99A plasmid expressing wild-type Tar as in FRET experiments (pSJAB21) to generate a high-variability strain. To maintain uniform growth conditions, the strain used as a wild type for swimming was transformed with an empty pTrc99A plasmid. Both high-variability and WT strains were transformed with a NaSal-inducible pRR31 plasmid expressing cytosolic EYFP to assist with wave tracking (pVS118, gift from Victor Sourjik).

For receptor ratio quantification on swim plates, an adaptation-proficient (CheRB+) version of TSS2155, TSS2144, was utilized, as adaptation is necessary for ascending gradients of attractants. We modulated the motility of TSS2144 by transforming the strain with an arabinose-inducible pBAD33 plasmid expressing WT FliC (pC100B-12, gift from Howard Berg). To determine the optimal levels of arabinose induction, we compared the motility of strain TSS2144/pC100B-12 to that of WT strain TSS2096 carrying an empty pBAD33 plasmid and ΔFliC strain TSS2097 carrying pC100B-12 on rich media soft agar swim plates (EDF 1c). The same strain, TSS2155, without the plasmid expressing WT FliC, was used as a non-motile sensor for gene expression.

Finally, wild-type MG1655 was also used to determine tumble bias and run speed at different growth rates in rich and minimal media.

For an overview of all the strains used in this study, refer to Table 1.

#### 2 Growth conditions

For experiments conducted in rich media, cells were retrieved from a -80°C glycerol-based stock and inoculated in 2 ml tryptone broth (TB; 1% bactotryptone, 0.5% NaCl, adjusted to pH 7.0) supplemented with 100 $\mu$g/mL ampicillin and 34 $\mu$g/mL chloramphenicol to maintain plasmids when necessary. The cultures were then incubated overnight at 30°C and 400 RPM until saturation. Subsequently, cells from the saturated overnight culture were diluted 1:50 (except for figure 4b, where the initial dilution was varied) in 10 ml of TB and supplemented with 100 $\mu$g/mL ampicillin and 34 $\mu$g/mL chloramphenicol, along

**Table 1** Strains used in this study.

| Background Strain | Background Strain Source | Background Strain Genotype | Plasmid 1 | Plasmid 2 |
|---|---|---|---|---|
| TSS1735 | [2] | MG1655<br>6TetO<br>6LacO<br>ΔLac | pOB2 | - |
| TSS2096 | V. Sourjik | WT MG1655 | - | - |
| TSS2096 | V. Sourjik | WT MG1655 | pTrc99A | - |
| TSS2096 | V. Sourjik | WT MG1655 | pBAD33 | - |
| TSS2096 | V. Sourjik | WT MG1655 | pSJAB21 | pVS118 |
| TSS2096 | V. Sourjik | WT MG1655 | pTrc99A | pVS118 |
| TSS2097 | V. Sourjik | MG1655 ΔFliC | - | - |
| TSS2097 | V. Sourjik | MG1655 ΔFliC | pC100B-12 | - |
| TSS2114 | This study | MG1655 ΔCheYZ | pSJAB106 | - |
| TSS2191 | This study | MG1655<br>FliC*<br>ΔCheRBYZ | pSJAB106 | - |
| TSS2191 | This study | MG1655<br>FliC*<br>ΔCheRBYZ | pSJAB21 | pSJAB196 |
| TSS2144 | This study | MG1655<br>ΔFliC::FLP<br>Tar-mCherry<br>Tsr-YFP | - | - |
| TSS2144 | This study | MG1655<br>ΔFliC::FLP<br>Tar-mCherry<br>Tsr-YFP | pC100B-12 | - |
| TSS2155 | This study | MG1655<br>ΔFliC::FLP<br>Tar-mCherry<br>Tsr-YFP<br>ΔcheRB | - | - |
| TSS2155 | This study | MG1655<br>ΔFliC::FLP<br>Tar-mCherry<br>Tsr-YFP<br>ΔCheRB | pMV13 | - |

with appropriate inducer concentrations to induce plasmids (refer to Table 2). For swimming competition experiments in serine gradients cells were grown 1:50 in 20 ml TB, to achieve higher cell counts within the bacterial wave. For supernatant experiments, cells were grown 1:50 in 50 ml TB, to collect a large volume of supernatant. All cultures were grown at 33.5°C with shaking at 250 RPM. Optical density (OD) was measured at 600 nm using a spectrophotometer (Genesys 10vis, Thermo Fisher Scientific), with a permitted error of ±0.01 for all reported OD values.

For experiments conducted in minimal media, cells were retrieved from the same -80°C glycerol-based stock and inoculated in 2 ml H1 minimal salts medium (MMH1; 50 mM KPO4, 0.5 mM MgSO4, 7.6 mM (NH4)2SO4, 1.25 $\mu$M Fe2(SO4)3, adjusted to pH 7.0) supplemented with 0.5% v/v glycerol, 0.01% w/v thiamine hydrochloride, and 100 $\mu$g/mL ampicillin when necessary to maintain plasmids. Cultures were then incubated for approximately 2 days at 30°C and 400 RPM until saturation. Subsequently, cells from the saturated culture were diluted at least 1:200 in 10 ml of MMH1 supplemented with the same supplements indicated before and appropriate inducer concentrations to induce plasmids (refer to Table 2). Cultures were grown overnight at 33.5 °C with shaking at 250 RPM. As in the experiments conducted in rich media, OD was measured at 600 nm, with a permitted error of ±0.01 for all reported OD values.

For receptor promoter experiments in minimal media, cells were incubated overnight in TB, as described above, and then they were washed with 2 ml MMH1 without glycerol or thiamine hydrochloride three times. Cells were then inoculated in fresh MMH1 supplemented with the indicated carbon source.

**Table 2** Plasmids used in this study. Amp: 100 $\mu$g/mL ampicillin; Cam: 34 $\mu$g/mL chloramphenicol.

| **Plasmid** | **Source** | **Product** | **System** | **Induction** | **Resistance** |
|---|---|---|---|---|---|
| pSJAB21 | This study | WT Tar | pTrc99A | 30 $\mu$M IPTG | amp |
| pMV13 | This study | Tar-mCherry | pTrc99A | 30 $\mu$M IPTG | amp |
| pSJAB106 | [1] | CheY-mRFP1<br>CheZ-mYFP | pTrc99A | 50 $\mu$M IPTG<br>(rich media)<br>15 $\mu$M IPTG<br>(minimal media) | amp |
| pSJAB196 | This study | CheY-mRFP1<br>CheZ-mYFP | pRZ30 | 8.5 $\mu$M NaSal | cam |
| pVS118 | V. Sourjik | EYFP | pRR31 | 10 $\mu$M NaSal<br>(aspartate waves)<br>20 $\mu$M NaSal<br>(serine waves) | cam |
| pC100B-12 | H. Berg | WT FliC | pBAD33 | 0.01% Arabinose | cam |
| pOB2 | [2] | LacI-mCherry<br>TetR-EYFP | pBAD24 | no induction<br>(relied on<br>promoter<br>leakiness) | amp |

**Table 3** Parameters used to fit the mixed-species MWC model

| **Parameter** | **Value** | **Reference** |
|---|---|---|
| C | 0.314 | [3] |
| $\epsilon_0$ | 0.826 | [3] |
| $\epsilon_A$ | 1.23 | [3] |
| $\epsilon_s$ | 1.54 | [3] |
| $\tilde{K}$ | 30 $\mu$M | - |
| $N_{\text{total}} = N_A + N_s$ | 32 (rich media)<br>100 (minimal media) | - |

### 3 Single-cell FRET microscopy

For single-cell FRET microscopy, cells (strain TSS2191/pSJAB106 for WT experiments or TSS2191/pSJAB21/pSJAB196 for high variability experiments) were collected by centrifugation (5 min at 5,000 RPM) and washed twice in 10mL motility media (MotM; 10 mM KPO4, 0.1 mM EDTA, 1 $\mu$M L-methionine, and 10 mM lactic acid, adjusted to pH 7.0). Following this, cells were resuspended in MotM and incubated at room temperature (approximately 22°C) for 90 minutes prior to imaging, to allow for further fluorophore maturation. In each experiment, cells immobilized on a glass coverslip were placed in a flow cell under continuous flow of MotM (400 $\mu$L/min) regulated by a syringe pump (PHD 2000, Harvard Apparatus). MotM solutions were utilized to incrementally add and remove varying concentrations of the attractants L-serine or L-aspartate (Sigma-Aldrich), with each concentration sustained for approximately 80 seconds.

Imaging was conducted using an inverted microscope (Eclipse Ti-E, Nikon) equipped with an oil-immersion 100x 1.45 NA phase-contrast objective lens (Nikon). Illumination occurred every 2 seconds using a 500 nm LED (pE-4000, CoolLED) with a pulse duration of 20 milliseconds for experiments with serine stimuli or a broad-spectrum LED (SOLA SE, Lumencor) with a pulse duration of 50 milliseconds for experiments with aspartate stimuli. For experiments with serine stimuli, epifluorescent light was split into two channels via a 2-camera image splitter (TwinCam, Cairn Research) equipped with a 580 nm dichroic mirror (Semrock) and two emission filters (520 nm and 593 nm, Semrock), each feeding into identical sCMOS cameras (ORCA-Flash 4.0 V2, Hamamatsu), capturing donor (YFP) and acceptor (RFP) channels separately. For experiments with aspartate stimuli, epifluorescent light was split into two channels via a single camera image splitter (OptoSplit II, Cairn Research) equipped with a 580 nm dichroic mirror (Semrock) and two emission filters (542 nm and 641 nm, Semrock), each projecting side-by-side into a single sCMOS camera (ORCA-Flash 4.0 V2, Hamamatsu), capturing donor (YFP) and acceptor (RFP) channels simultaneously.

FRET images were binned by a factor of 4x4 pixels to maximize their signal-to-noise ratio. All experiments were conducted at room temperature (approximately 22°C). FRET dose-response experiments were highly reproducible (EDF 2). Control experiments verified negligible growth throughout the entire duration of the experiment (EDF 7).

### 4 FRET image analysis

Image analysis was conducted using custom MATLAB scripts. Initially, to compensate for any movement of the flow cell, all images were registered based on the first image of the experiment by employing a rigid transformation. Subsequently, alignment of donor and acceptor channels was achieved through an affine transformation. Following alignment, cell segmentation was performed on the donor (YFP) channel (given its typically higher brightness compared to the acceptor channel), utilizing a modified Otsu algorithm. After segmentation, a rectangular region of interest (ROI) of constant area was defined for each cell. The average YFP and RFP intensities over the ROI were then extracted as a function of time. To mitigate the drift in fluorescence intensities primarily attributed to fluorophore bleaching, the FRET ratio of each cell (defined as RFP/YFP) was fitted with a double exponential function. Subsequently, the FRET ratio was divided by the double exponential and normalized between one (unstimulated steady-state activity) and zero (activity upon addition of a saturating dose of the attractant serine or aspartate), which corresponds to the kinases CheA in the cell being fully active and fully inactive, respectively. Therefore, the kinase activity of each cell is given by:

$$\alpha(t) = \frac{FRET(t) - FRET_{\text{saturating}}}{FRET_{\text{steady-state}} - FRET_{\text{saturating}}}$$

### 5 MWC Model

To obtain a distribution of $K_{1/2}$ parameters from the receptor expression data, we employed the mixed-species MWC model as described in [3]. The expression for the normalized response to L-serine is:

$$a = \frac{\epsilon_0 \epsilon_s^{N_s} \epsilon_A^{N_A} \left(1 + C\frac{[L]}{\tilde{K}}\right)^{N_s}}{\left(1 + \frac{[L]}{\tilde{K}}\right)^{N_s} + \epsilon_0 \epsilon_s^{N_s} \epsilon_A^{N_A} \left(1 + C\frac{[L]}{\tilde{K}}\right)^{N_s}}$$

where $\epsilon_A$, $\epsilon_s$, and $\epsilon_0$ are the L-serine binding energies to Tar, Tsr, and to the three minor receptors, respectively. The number of Tsr receptors is denoted by $N_s$, while the number of Tar receptors is denoted by $N_A$. The vector $[L]$ represents the concentrations of L-serine and $C$ and $\tilde{K}$ describe the ligand dissociation constant for the active state of the receptor, as:

$$K_A = \frac{\tilde{K}}{C}$$

All variables apart from parameters $N_s$ and $N_A$ were kept constant for all cells and are given in Table 3. The L-serine concentration vector $[L]$ was defined as a logarithmically spaced vector between 0.1 and 10,000. To transform the measured Tar/Tsr ratio to the MWC receptor count $N_s$ and $N_A$, we used:

$$N_s = \frac{N_{\text{total}}}{\frac{\text{Tar}}{\text{Tsr}} + 1}$$

$$N_A = N_{\text{total}} - N_s$$

The normalized expression for the response to L-serine yielded a sigmoidal function between 0 and 1 for all measured receptor ratios in rich media. For minimal media experiments, a fraction of cells with extreme receptor ratios did not produce a response between 0 and 1. These data points were excluded from $K_{1/2}$ extraction. It is likely that these cells correspond to the fraction of non-motile cells, which is known to be larger for cells grown in minimal media compared to cells grown in rich media [4]. Finally, to extract $K_{1/2}$ from the MWC normalized response to L-serine we used a Hill function of the form:

$$a = \frac{1}{1 + \left(\frac{[L]}{K_{1/2}}\right)^H}$$

where $K_{1/2}$ and $H$ are the fit parameters. The same Hill function was used to fit experimental dose-response data.

### 6 Colony expansion rate quantification

To calibrate the FRET pair (pSJAB106) and WT FliC plasmid expression (pC100B-12), we assessed the expansion rate of bacterial colonies carrying the plasmid of interest (strain TSS2114/pSJAB106 or TSS2097/pC100B-12) alongside WT colonies (strain TSS2096) carrying an empty version of the same plasmid (EDF 1). Cells grown in rich media were collected at OD = 0.45 and cells grown in minimal media were collected at OD = 0.30 and diluted to OD = 0.01 using fresh media. Then, 10 $\mu$L of cell culture was inoculated in the center of semi-solid agar plates. Liquid media and agar plates were supplemented with appropriate antibiotics and inducers, refer to Table 2. All colony expansion rate experiments were conducted at 33.5°C and 100% humidity. For each experiment, six swim plates were imaged simultaneously using a custom-built motorized turret. Images of each plate were taken every 10 minutes using a Canon DSLR camera. To extract the expansion rate, the diameter of the expanding colony was determined as a function of time by fitting an ellipse with an eccentricity smaller than 0.2 using the MATLAB regionprops function on every image, after subtracting the first image of each plate (taken before the colony had formed).

### 7 Chemoreceptor copy number quantification

The quantification of Tar and Tsr chemoreceptor copy numbers was conducted using both a plate reader assay and single-cell fluorescence microscopy. For growth and chemoreceptor copy number measurements in the plate reader (Victor X3 2030 Multilabel reader, PerkinElmer), cells (strain TSS2155) were diluted from an overnight saturated culture into fresh media (dilution ratios 1:50 for TB and 1:200 for MMH1) and 200$\mu$L of cell culture was aliquoted into each well of a 96-well plate (96-well Clear Flat Bottom TC-treated Microplate, Corning). Cultures were grown at 33.5 °C with double orbital shaking with a shaking amplitude of 2 mm until they reached saturation. Fluorescence (YFP and mCherry) and optical density (OD600) were measured every 15 minutes. The autofluorescence of the non-fluorescent parent strain of TSS2155, TSS2097, grown in the same well plate, was subtracted from all fluorescent measurements. The OD units of the plate reader were calibrated against the OD units of the spectrophotometer used to measure the OD of the batch cultures for single-cell fluorescence microscopy experiments (Genesys 10vis, Thermo Fisher Scientific) by constructing a calibration curve through serial dilution of a saturated culture. The receptor ratio units measured with the plate reader were calibrated against mean receptor ratios obtained from fluorescence microscopy for the same optical densities.

For single-cell quantification of chemoreceptor copy numbers, cells (strain TSS2155 for WT experiments or TSS2155/pMV13 for high variability experiments) were harvested from batch TB or MMH1 culture at the appropriate OD by centrifugation (5 min at 5,000 RPM). The cells were then washed twice in 10mL minimal motility media (minimal MotM; 10 mM KPO4 and 0.1mM EDTA, adjusted to pH 7.0). After washing, cells were diluted in minimal motility media, allowed to mature, and plated on agarose pads (1.5% agarose in minimal MotM). The pads were left to dry for 10 minutes before immediate imaging.

Imaging was performed using an inverted microscope (Eclipse Ti-E, Nikon) equipped with an oil-immersion 100x 1.45 NA phase-contrast objective lens (Nikon). YFP and mCherry fluorescence was excited using an LED system (pE-4000, CoolLED), and emissions were captured sequentially using a multi-band filter (Semrock) and directed into two identical sCMOS cameras (ORCA-Flash 4.0 V2, Hamamatsu) via a 2-camera image splitter (TwinCam, Cairn) equipped with a 580 nm dichroic mirror (Semrock). Approximately 20 fields of view, totaling around 1000 cells, were typically imaged per experiment. Measurements took place at room temperature (approximately 22°C). A control experiment revealed that nearly zero growth took place during the total duration of the experiment (EDF 7b).

To ensure accurate determination of the Tar/Tsr ratio, excitation pulse durations and intensities for YFP and mCherry were calibrated using a two-color fluorescent repressor-operator system (FROS) standard [2]. This FROS strain (strain TSS1735/pOB2) contained six mCherry- and YFP-repressor fusions on its chromosome. By using a 100 ms 500 nm LED excitation pulse for mYFP and a 200 ms 580 nm LED excitation pulse for mCherry (both at maximum intensity), nearly identical emission intensities for mCherry and YFP were achieved, effectively equating the measured mCherry/YFP fluorescent intensity ratio to the true Tar/Tsr receptor abundance ratio (EDF 3). To avoid movement of the chromosome due to diffusion during the measurement, cells were chemically fixed by incubating them for 10 min at room temperature in a 4% paraformaldehyde solution (Sigma-Aldrich).

### 8 Chemoreceptor image analysis

In each field of view, a set of images comprising one phase-contrast, one YFP, and one mCherry channel image was obtained. Alignment of images from the two cameras was performed using a custom MATLAB script through an affine transformation. Subsequently, cell segmentation was conducted on the phase-contrast channel utilizing a modified Otsu algorithm. We used the phase-contrast channel to segment the cells instead of one of the two fluorescent channels to ensure that cells with low fluorescence (and hence low receptor expression) are included in the analysis. Following segmentation, background subtraction was applied to each image, and a rectangular ROI of the same area is defined for each cell. The average mCherry and YFP fluorescent intensities over the ROI are extracted for every cell. When indicated, average fluorescent intensities are normalized by cell area, determined by fitting an ellipse on the detected cells in the phase-contrast channel

### 9 Fabrication of microfluidic chemostats

Microfluidic chemostats, commonly referred to as "mother machines," were fabricated using the silicone polymer polydimethylsiloxane (PDMS). The master mold for the PDMS device is a silicon wafer on which features are imprinted on a layer of SU-8, an epoxy-based negative photoresist [5]. Features are imprinted using UV light exposure through two separate photomasks. The first layer of the device comprises an array of shallow and narrow growth channels measuring 25 $\mu$m in length and 0.96 $\mu$m in height. To allow the tight confinement of cells of different widths and hence accommodate growth at different growth rates, the width of the growth channels varies within the range of 0.7 $\mu$m to 1.2 $\mu$m, with a step size of 0.1 $\mu$m. The edges of the channels were smoothed to reduce optical aberrations during phase-contrast imaging [6]. The second layer of the device features a deeper and wider feeding channel with low fluidic resistance, enabling rapid liquid exchange. The feeding channel measures 12 mm in length, 100 $\mu$m in width, and 32.5 $\mu$m in height. The designs for the two different layers were created using CAD software (klayout) and transferred onto two separate quartz photomasks (Photronics). Fabrication of the silicon master mold was conducted at AMOLF's cleanroom facilities. Following fabrication, a hydrophobic coat of chlorotrimethylsilane (Sigma-Aldrich) was applied via vapor deposition.

To cast PDMS devices, the master mold was coated with a layer of degassed 10:1 PDMS:curing agent mixture (Sylgard 184, Dow Corning) and cured at 80°C for 1 hour. After cooling the wafer to room temperature, individual devices were cut and separated from the wafer, and inlets and outlets for each feeding channel were created using a 0.75-mm biopsy punch (World Precision Instruments). To remove any residual uncured PDMS, devices were immersed in a pentane bath for 2 hours and then in an acetone bath twice for 2 hours each. The PDMS devices were then allowed to dry at room temperature overnight. To facilitate bonding, the PDMS devices were cleaned with transparent adhesive tape (Magic Tape, Scotch) and treated, along with glass-bottom dishes (GWST-5040, WillCo Wells), in a plasma cleaner (PDC-002, Harrick Plasma). Following a 60-second exposure to ambient air plasma under a 300 mTorr vacuum, the devices were laminated onto the glass-bottom dishes and baked on an 80°C hot plate for 5 minutes to establish a covalent bond. In order to passivate the PDMS surface and prevent cell attachment, the devices were incubated with a 10 mg/ml BSA solution for 45 minutes at 37°C to allow the BSA solution to be drawn into the growth channels via evaporation through the gas-permeable PDMS. The devices were then immediately used for experiments.

### 10 Supernatant experiments

For supernatant experiments, cells (TSS2155) grown in rich media were collected at OD = 0.30 and OD = 0.80 by centrifugation (5 min at 5,000 RPM) and washed twice in 10mL minimal MotM. The supernatant of those cultures was filtered using a 0.22 $\mu$m syringe filter and stored at 3°C until the experiment. The washed cells were imaged following the same protocol as for all receptor-labeled cells, with the exception that imaging took place at 33.5°C, so the measured receptor ratios are directly comparable to those of growing cells in the microfluidic chemostat.

Simultaneously, 10$\mu$L of the original OD = 0.30 culture was injected into a microfluidic chemostat through its inlet. The inlet and outlet of the chemostat were then sealed with adhesive tape (Magic Tape, Scotch) to prevent flow and the entire device was centrifuged for 20 minutes at 800 RPM, orienting the growth channels parallel to the centrifugal force axis to force cells into the growth channels. Following centrifugation, the device was incubated for 2 hours at 33.5°C, to allow for additional cells to enter the growth channels via diffusion. Subsequently, the collected supernatants were placed inside sterile glass reservoirs pressurized to 130 kPa using nitrogen. These reservoirs were then connected through separate tubes to a distribution valve (M Switch, Fluigent), which in turn was connected to the microfluidic device through a tube plugged into its inlet. The distribution valve input, and hence the growth environment of the cells, was controlled through a custom MATLAB script.

To extract the receptor ratio of the growing cells, the microfluidic device was imaged using the same microscopy setup and imaging settings we employed for all receptor-labeled cell experiments. However, imaging occurred at 33.5°C, instead of room temperature, to facilitate cell growth. To extract the morphology of the cells and to determine a segmentation mask, phase-contrast images were acquired from a single field of view every minute. Furthermore, at 20-minute intervals, corresponding to the maximum division rate observed in our experiment, one mCherry and one YFP image were acquired to quantify receptor expression.

### 11 Image analysis of microfluidic chemostat experiments

To accurately segment bacteria cells growing in the microfluidic chemostat, we trained a machine learning model to predict the outline of each cell. We used a convolutional neural network (CNN) based on the U-Net architecture [7, 8] to transform the original microscopy images into maps of cell outlines using custom Python scripts. To train the network, we simultaneously acquired fluorescent images, using the bright CheY-mRFP1 fusion, and phase-contrast images of growing cells. Subsequently, we manually created binary masks of all cell outlines in 500 CheY-mRFP1 512x512 pixel images and augmented this dataset using geometric transformations such as scaling, translation, and rotation. This augmented dataset was then utilized to train the CNN.

Following the training phase, we applied the CNN to transform 2000 CheY-mRFP1 frames, distinct from the original training dataset, without any post-processing of the images. The CNN generated unique 512x512 pixel output matrices for each image, with high scores corresponding to cell outlines while cell bodies and the background contained nearly zero scores. We then re-trained the CNN using the phase-contrast images and the binary masks created by the CNN based on CheY-mRFP1 images as inputs. Additionally, we augmented these datasets further using geometric transformations. Through this training method, we increased our training dataset by a factor of 4, while manually segmenting only CheY-mRFP1 images, which are substantially easier to segment compared to phase-contrast images due to the high brightness of the fluorescent fusion protein and the near-zero background fluorescence.

The primary advantage of using phase-contrast images for cell segmentation is the elimination of the need for an additional fluorescent protein and imaging channel, as well as the avoidance of phototoxicity associated with fluorescent imaging. Visual inspection of the transformed images revealed consistent cell masks produced by the CNN, irrespective of whether the original image was obtained through fluorescence or phase-contrast microscopy.

Subsequently, the CNN output binary matrices underwent further processing by thresholding using a modified Otsu algorithm and removing small clusters of unconnected pixels, effectively setting only the pixels corresponding to cell outlines to unity and every other pixel to zero. Then, alignment of images from the two cameras was performed using a custom MATLAB script employing an affine transformation. All images were then registered based on the first image of the experiment using a rigid transformation to correct for movement of the microfluidic device. Fiducial markers in the form of growth channel numbering were incorporated into the device to assist with image registration. Finally, individual growth channels were cropped out by summing the fluorescence of both channels and detecting peaks in the fluorescence intensity profile.

For cell segmentation, we employed the MATLAB regionprops function, which detects cells in the CNN-transformed images as elliptical objects. For each cell, we determined its minor and major axes, along with the x- and y-coordinates of its centroid. The centroid was then tracked across all frames, under the assumption that interframe movement could not exceed a threshold defined as one quarter of the shortest cell length in any given image. Movement was considered only along the long axis of the growth channel, as the cells were constrained along their width, thereby minimizing tracking to a one-dimensional problem. Divisions were identified by abrupt changes in cell length (the major axis of the fitted ellipse), coupled with an increase in cell count. Conversely, a reduction in cell count, alongside the disappearance of centroids near the origin of the growth channel, indicated cells exiting towards the feeding channel. Subsequently, the minor and major axes of the fitted ellipses were utilized to define rectangular segmentation masks, facilitating the extraction of fluorescent intensity for each cell across all frames. Areas devoid of cells were used to extract the background intensity of each frame.

### 12 Fabrication of microfluidic devices for swimming competition experiments

For swimming competition experiments, we utilized a microfluidic device comprising a long linear channel with dimensions of 1.2 mm x 3 cm x 100 $\mu$m (width, length, height). These devices were fabricated following the procedure outlined in [9]. To establish a cost-effective method of producing devices without the necessity for silicon wafer masters, we manufactured epoxy molds utilizing the silicon wafer-molded PDMS devices as masters [10]. The devices were submerged in a degassed mixture of epoxy and hardener (epoxacast 690, Smooth-On) in a 10:3 ratio, and the mixture was allowed to solidify at room temperature for 2 days. Similar to silicon wafers, a hydrophobic coat of chlorotrimethylsilane was applied via vapor deposition.

To cast PDMS devices, the master mold was coated with a layer of degassed 10:1 PDMS:curing agent mixture (Sylgard 184, Dow Corning) and cured at room temperature for 2 days due to the low glass transition temperature of the epoxy mold. Individual devices were then cut and separated from the wafer, and a 0.75-mm outlet and a 2-mm cell reservoir were created on either side of the linear channel using biopsy punches (World Precision Instruments). Subsequently, the devices underwent cleaning with transparent adhesive tape (Magic Tape, Scotch), followed by rinsing sequentially with isopropanol, methanol, and Millipore-filtered water. After drying the devices using nitrogen, they were treated, alongside glass-bottom dishes (GWST-5040, WillCo Wells), in a plasma cleaner (PDC-002, Harrick Plasma). Following a 60-second exposure to ambient air plasma under a 300 mTorr vacuum, the devices were laminated onto the glass-bottom dishes and baked on an 80°C hot plate for 5 minutes to establish a covalent bond. Then, after cooling the devices to room temperature, they were immediately utilized for experiments.

### 13 Swimming competition experiments

For swimming competition experiments, cells (TSS2096/pTrc99A/pVS118 for WT experiments or TSS2096/ pSJAB21/pVS118 for high-variability experiments) grown in rich media were collected at the appropriate OD by centrifugation (5 min at 5,000 RPM) and washed twice in 10mL MotM. Cells were then concentrated to OD = 1.0 for aspartate wave experiments or OD = 2.0 for serine wave experiments. In parallel, the microfluidic linear channel devices were filled with either 100$\mu$M L-aspartate or 100$\mu$M L-serine (Sigma-Aldrich) in MotM supplemented with 0.05% w/v polyvinylpyrrolidone-40 (Sigma-Aldrich) from the 0.75-mm outlet. Subsequently, excess liquid was removed from the 2-mm cell reservoir and 5$\mu$L cell suspension was loaded gently into the reservoir. The outlet was sealed with adhesive tape (Magic Tape, Scotch) to prevent flow due to differences in hydrostatic pressure across the device.

Migrating waves in swimming competition experiments were imaged immediately after cell loading using an inverted microscope (Eclipse Ti-E, Nikon) equipped with an environmental chamber (Okolab) set to a temperature of 33.5°C and 80% humidity. YFP images of the bacterial wave were captured through a 10x 0.30 NA objective lens (Nikon). The field of view was illuminated with a 500 nm LED (pE-4000, CoolLED) set to maximum intensity, with an exposure time of 100 ms. Images were recorded by a sCMOS camera (ORCA-Flash 4.0 V2, Hamamatsu). To track the bacterial wave, the microscope stage (MicroStage, Mad City Labs) was programmed using a custom MATLAB script to translocate every 10 seconds by one frame width (1.33 mm). Six consecutive images were taken for each sweep, and full sweeps occurred every 60 seconds until the wave had traversed all six consecutive fields of view. A control experiment revealed that nearly zero growth took place during the total duration of the experiment (EDF 7c and EDF 7d).

For wave speed analysis, raw YFP images were stitched into single images for each sweep using custom MATLAB scripts. The initial stitched image, which lacked cells, was subtracted from subsequent images to eliminate imaging artifacts. Images were then segmented using a modified Otsu algorithm, filtering out objects smaller than 100 pixels. Subsequently, image intensities were normalized between one and zero, and a Gaussian function was fitted to the intensity profile along the long axis of the channel (EDF 6). The wave position was determined by identifying the peak of each Gaussian, and velocity was calculated by extracting the distance between successive peaks and dividing it by the time between sweeps (60 seconds).

### 14 Chemoreceptor copy number quantification on swim plates

Experiments with swim plates were performed similarly to what we reported on [11].

Briefly, to quantify the chemoreceptor copy number of cells forming colonies on swim plates, cells (strain TSS2144/pC100B-12) grown in rich media were collected at OD = 0.45 and diluted to OD = 0.01 using fresh media. Then, 10 $\mu$L of cell culture was inoculated in the center of semi-solid agar plates. The agar plates were made using rich media, as described above, supplemented with 0.26% bacteriological agar (Avantor). Liquid media and agar plates were supplemented with appropriate antibiotics and inducers, refer to Table 2. The plates were incubated at a temperature of 33.5°C and 100% humidity until the colonies reached the edge of the plate. Cells from the center and the edge of the migrating colony were collected using pipette tips and diluted into minimal MotM for subsequent fluorescent imaging. To quantify chemoreceptor numbers, we followed the same fluorescent imaging protocol as for receptor-labeled cells grown in liquid cultures.

For experiments involving non-motile cells used as biosensors for gene expression, cells (strain TSS2144) grown in rich media were collected at OD = 0.45 and mixed with fresh rich media supplemented with 0.26% bacteriological agar to achieve a final OD = 0.01. Swim plates were allowed to solidify at room temperature for 2-3 hours, following which 10 $\mu$L of OD = 0.01 motile cell culture (strain TSS2096) was inoculated in the center of the semi-solid agar plates. The motile population of cells shapes the gradient of the plate, ensuring that the non-motile biosensor strain encounters the same local environment as motile cells. Once the migrating cell colony reached the edge of the plate, cells from the center and the edge of the colony were collected using pipette tips and diluted into minimal MotM for subsequent fluorescent imaging.

### 15 Fabrication of microfluidic devices for single cell swim tracking

To track the run-and-tumble motion of single cells, we utilized a microfluidic device comprising a long linear channel with dimensions of 1.2 mm x 3 cm x 60 $\mu$m (width, length, height). These devices were fabricated following the procedure outlined in [11]. Briefly, the master mold was coated with a layer of degassed 10:1 PDMS:curing agent mixture (Sylgard 184, Dow Corning) and cured at 80°C for 12 hours. After cooling the wafer to room temperature, individual devices were cut and separated from the wafer, and a 0.75-mm outlet and a 0.75-mm inlet were created on either side of the linear channel using biopsy punches (World Precision Instruments). Subsequently, the devices underwent cleaning with transparent adhesive tape (Magic Tape, Scotch), followed by rinsing sequentially with isopropanol, methanol, and Millipore-filtered water. A 22 mm x 50 mm glass coverslip was rinsed sequentially with acetone, isopropanol, methanol, and Millipore-filtered water. After drying the devices and the coverslips using nitrogen, they were treated in a plasma cleaner (PDC-32G, Harrick Plasma). Following a 60-second exposure to ambient air plasma under a 300 mTorr vacuum, the devices were laminated onto the glass coverslips and baked on an 80°C hot plate for 5 minutes to establish a covalent bond. Then, after cooling the devices to room temperature, they were immediately utilized for experiments.

### 16 Swim tracking experiments

For swim tracking experiments, WT cells (strain TSS2096) grown in rich or minimal media were collected at the appropriate OD by centrifugation (5 min at 5,000 RPM) and washed twice in 10mL MotM supplemented with 0.05% w/v polyvinylpyrrolidone-40 (Sigma-Aldrich). Cells were then diluted to an OD of 0.001. Subsequently, 10 $\mu$L of the mixture was introduced into the device from the inlet and excess liquid was removed from the outlet. Both the outlet and the inlet were then sealed with adhesive tape (Magic Tape, Scotch) to prevent flow due to differences in hydrostatic pressure across the device.

Swimming cells were imaged immediately after cell loading using an inverted microscope (Eclipse Ti-E, Nikon) equipped with a custom environmental chamber set to a temperature of 30°C and 50% humidity for experiments in Figures 5e and EDF 9 or 33.5°C and 80% humidity for experiments in EDF 6c and EDF 6d, to match the environmental conditions of swimming competition experiments. Phase-contrast images of swimming bacteria were captured through a 4x 0.13 NA phase objective lens (Nikon). Images were recorded by a sCMOS camera (ORCA-Flash 4.0 V2, Hamamatsu) at 20 frames per second.

To detect and track cells and extract their tumble bias and run speed, a custom MATLAB script was used as described in [12]. Trajectories shorter than 10 seconds were excluded for further analysis. In a typical experiment, 4 fields of view where imaged for 200 seconds each, containing a total of at least 1000 cell trajectories.

### 17 Growth rate measurements

To assess the growth rate of the FRET (TSS2191/pSJAB106), receptor-labeled (TSS2155), and WT (TSS2097) strains, we employed a microplate reader (Epoch 2 microplate reader, BioTek). Overnight cultures in rich media were diluted 1:50 into fresh media and overnight cultures in minimal media were diluted 1:200 into fresh media. For the FRET strain, the appropriate antibiotic and inducer concentrations were added to the media (refer to Table 2). Subsequently, 200 $\mu$L of diluted cells were aliquoted into each well of a 96-well plate (Costar 3595, Corning) and loaded into the plate reader. The plates were incubated at 33°C while continuously shaking in a double orbital pattern (1 mm radius, 807 cpm). The OD of each well was measured every 10 minutes for 12 hours for rich media and every 20 minutes for 24 hours for minimal media. The growth rates measured in the plate reader were within 10% of those obtained by sampling the shaking batch culture used for microscopy experiments.

### 18 Growth rate quantification

The quantification of *tar* and *tsr* promoter activities as a function of growth rate was conducted using a plate reader assay, as described above. The growth rate of the bacteria is expressed as doublings per hour and is calculated as:

$$\text{growth rate} = \frac{\log_{10}(\text{OD}_{t_2}) - \log_{10}(\text{OD}_{t_1})}{(t_2 - t_1)\log_{10}(2)}$$

### 19 Measurements of amino acid concentrations

L-aspartate and L-serine concentrations were quantified in cultures of the FRET strain (TSS2191/pSJAB106) grown using the standard protocol described for FRET experiments. At defined cell density (OD) intervals, 200 $\mu$L samples were collected from the same growing culture and immediately placed on ice. After completing the time course, samples were thawed at room temperature and cells were separated from the supernatant by centrifugation.

L-aspartate and L-serine concentrations in the supernatant were measured using commercial colorimetric assay kits (L-aspartate: ab102512, Abcam; L-serine: ab241027, Abcam), following the manufacturer's instructions. To ensure measurements fell within the linear detection range of each assay, 10 $\mu$L samples were diluted 1:5 in aspartate assay buffer for L-aspartate quantification and 1:6 in serine assay buffer for L-serine quantification.

For both assays, absorbance was measured at 570 nm in 96-well plates using a microplate reader (Epoch 2, BioTek). Prior to measurement, plates were degassed in a vacuum desiccator to remove air bubbles, which can otherwise introduce significant artifacts in optical density readings.

For the L-aspartate assay, aspartate is enzymatically converted to pyruvate, which is subsequently oxidized to generate a colored product. Because pyruvate is present in the growth medium (at approximately 10% of the L-aspartate concentration), each sample was measured both in the presence and absence of the pyruvate converter enzyme. The signal obtained without the converter enzyme was subtracted to correct for background pyruvate.

For L-serine quantification, the assay detects D-serine. Therefore, each sample was measured both in the presence and absence of the serine racemase enzyme, which converts L-serine to D-serine. The L-serine concentration was calculated as the difference between total serine (measured with racemase; D+L) and D-serine (measured without racemase).

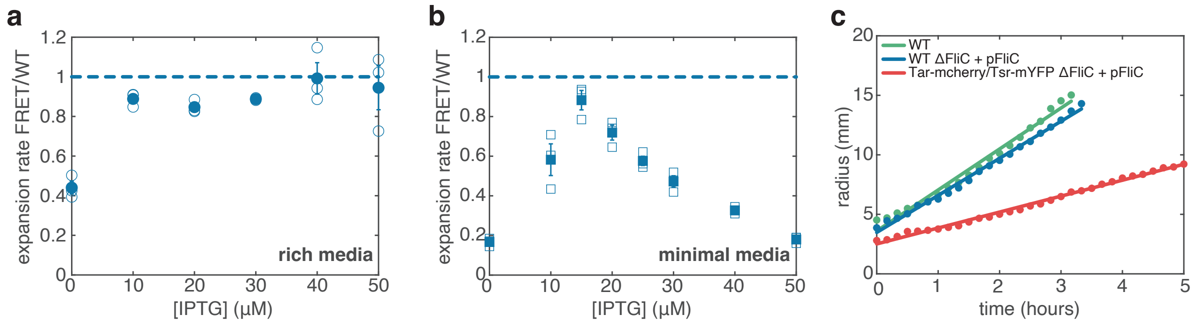


**EDF 1** Expansion rate of FRET and receptor-labeled strains. **(a)** Cells expressing CheY and CheZ protein fusions (TSS2114/pSJAB106), used for FRET experiments, yield chemotactic phenotypes with expansion rates nearly identical to WT cells (TSS2096/pTrc99A) when inoculated on rich media soft agar swim plates and induced with 40 or 50 $\mu$M IPTG. Hollow symbols represent technical replicates (three for each inducer concentration), while filled symbols represent the mean values of the three replicates. Error bars indicate the standard error of the mean and are typically the size of the graph points. **(b)** Similar to panel a, but for minimal media soft agar swim plates. When induced with 15 $\mu$M IPTG, the FRET strain yields an expansion rate nearly identical to that of WT cells. **(c)** Colony expansion of receptor-labeled strain as a function of time. Tar and Tsr protein fusions, used in receptor quantification experiments, produce chemotactic phenotypes when inoculated on rich media soft agar swim plates. Lines represent linear fits to the data points to determine the expansion rates. The expansion rates for the different strains are as follows: 3.23 $\pm$ 0.22 mm/hour (WT), 3.10 $\pm$ 0.20 mm/hour (WT $\Delta$FliC + pFliC), and 1.28 $\pm$ 0.06 mm/hour (Tar-mCherry/Tsr-YFP $\Delta$FliC + pFliC). The abbreviation pFliC denotes pBAD33-based plasmid pC100B-12 expressing WT FliC. WT carries empty pBAD33 (TSS2096/pBAD33), WT $\Delta$FliC + pFliC is strain TSS2097/pC100B-12, and Tar-mCherry/Tsr-YFP $\Delta$FliC + pFliC is strain TSS2144/ pC100B-12. Averages and standard errors of the mean are derived from two technical replicates.

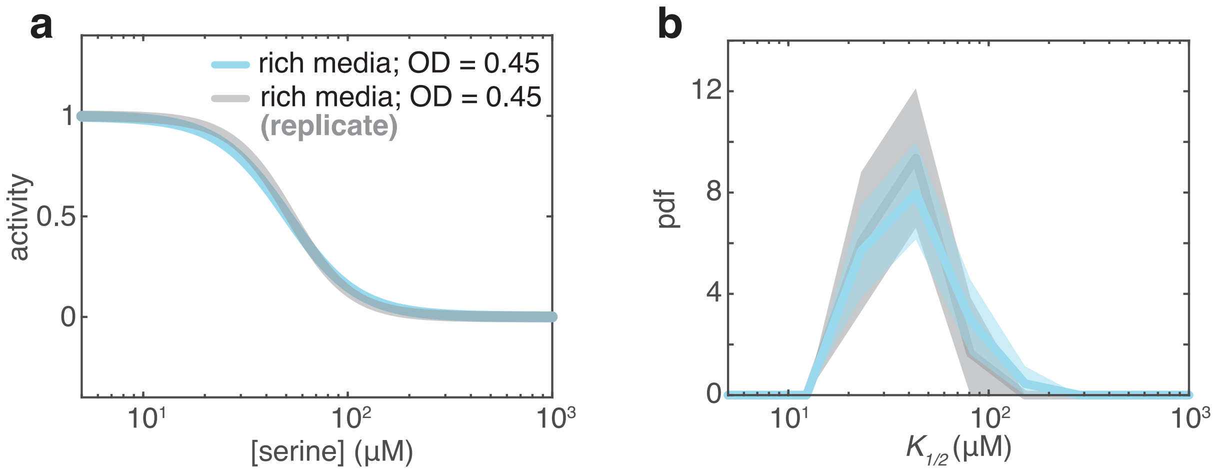


**EDF 2** Reproducibility of FRET dose-response experiment. **(a)** Hill function fits to the population-averaged L-serine dose-response data for cells grown to OD = 0.45 in rich media. The blue dose-response curve is replotted from Figure 1b. The gray dose-response curve represents a biological replicate performed on a different day with different ligand dilutions and is identical to the original dose-response curve. **(b)** Histograms of $k_{1/2}$ values obtained from single-cell dose-response curves for cells grown to OD = 0.45 in rich media. The blue distribution is replotted from Figure 1b, and the gray distribution is extracted from single-cell dose-response curves from the same biological replicate as in panel a, showing identical of $k_{1/2}$ distributions. Shaded areas represent 95% confidence intervals obtained through bootstrap resampling of the data.

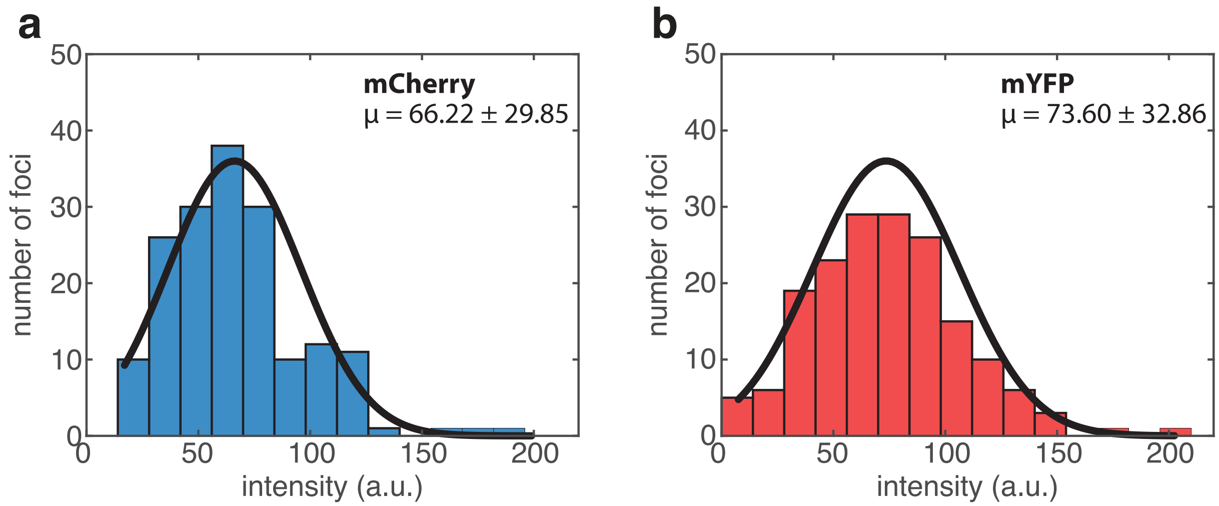


**EDF 3** Equating the Tar-mCherry/Tsr-mYFP fluorescence intensity ratio to Tar/Tsr receptor abundance ratio. A chemically fixed FROS strain containing six mCherry- and mYFP-repressor fusions on its chromosome (panels **a** and **b**, respectively) was imaged using the same settings and optical path as used in receptor-labeled experiments. The mean measured mCherry/mYFP fluorescence intensity ratio ($\mu$ denotes mean value $\pm$ standard deviation) was close to unity, indicating that the Tar-mCherry/Tsr-mYFP fluorescence intensity ratio effectively reflects the Tar/Tsr receptor abundance ratio.

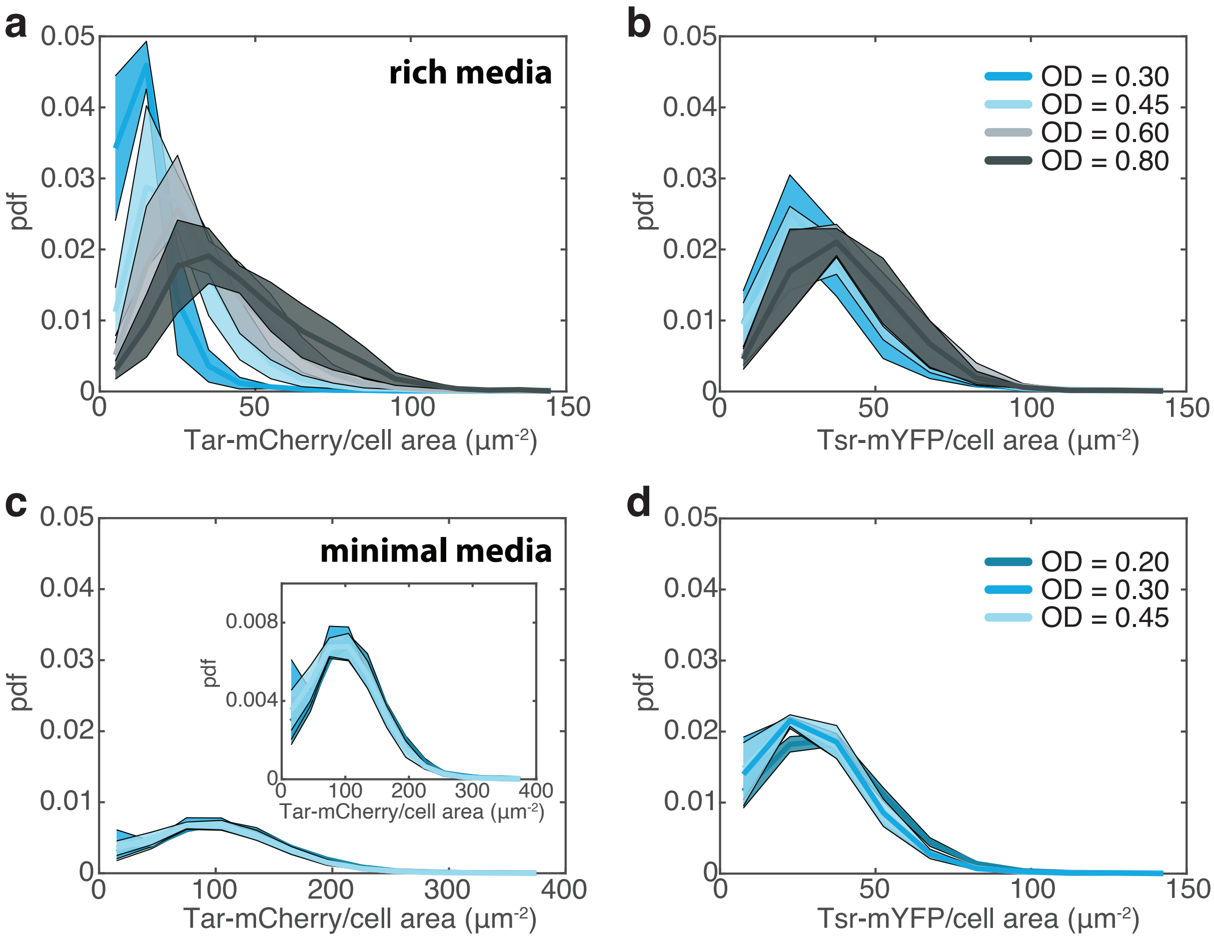


**EDF 4** Tar and Tsr chemoreceptor expression under different growth conditions. **(a)** Histograms of Tar-mCherry expression measured with fluorescence microscopy for cells grown to different optical densities (ODs) in rich media. Solid lines represent the mean of four independent biological replicates, with shaded areas indicating the standard error of the mean. The Figure legend and color scheme match panel b. **(b)** Same as panel a, but for Tsr-mYFP. **(c)** Histograms of Tar-mCherry expression for cells grown to different ODs in minimal media. Note the extended x-axis range. Solid lines represent the mean of three independent biological replicates, with shaded areas indicating the standard error of the mean. The Figure legend and color scheme match panel d. Inset: expanded version of the plot. **(d)** Same as panel c, but for Tsr-mYFP.

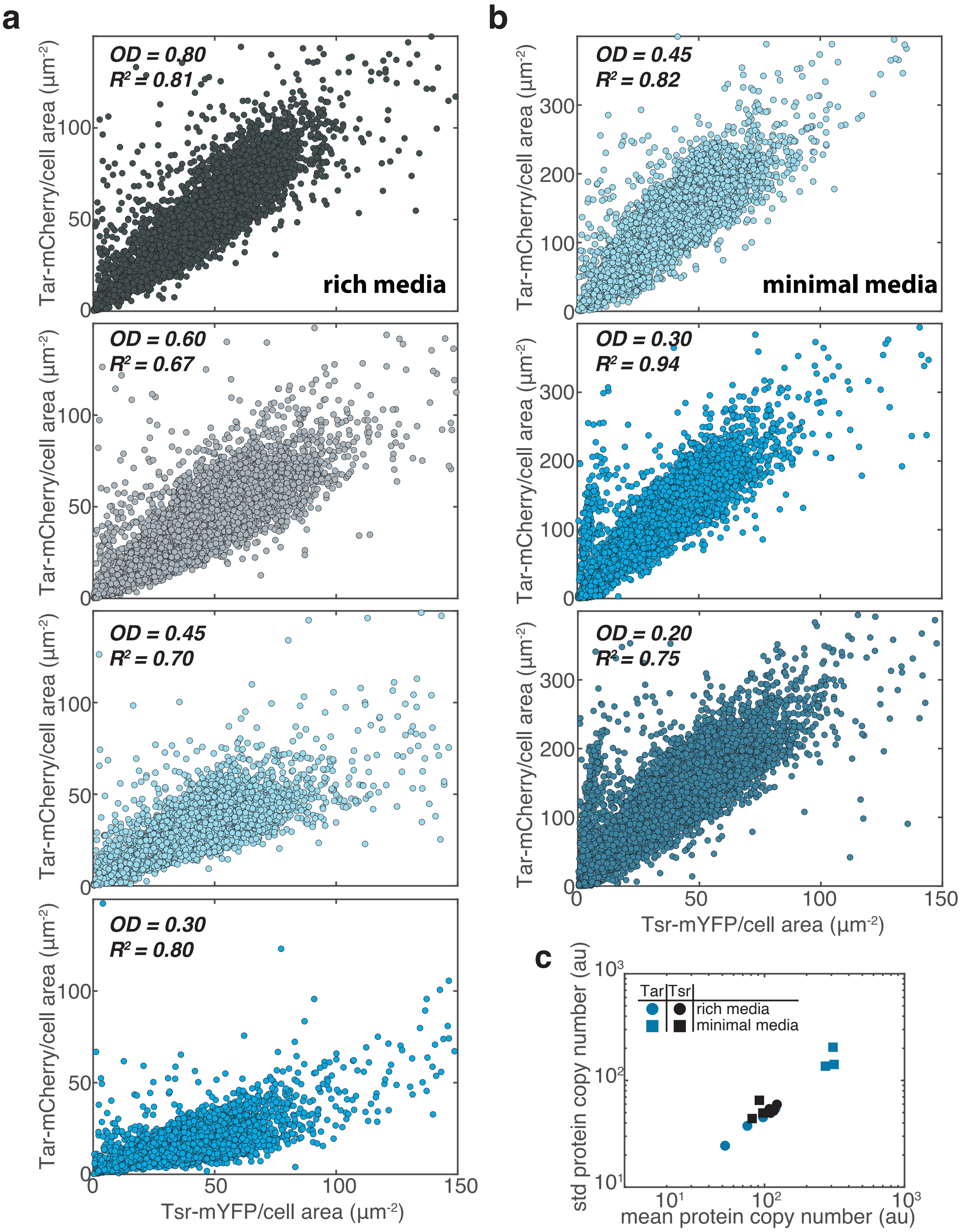


**EDF 5** Tar and Tsr chemoreceptor expression is corelated under all growth conditions. **(a)** Correlation between Tar-mCherry and Tsr-mYFP expression for cells grown to different optical densities (ODs) in rich media, measured using fluorescence microscopy. Each point represents a single cell, derived from four independent biological replicates. **(b)** Similar to panel a, but for cells grown in minimal media, with points obtained from three independent biological replicates. **(c)** Scaling of the mean and standard deviation of protein copy numbers (in arbitrary units) for Tar and Tsr across all conditions.

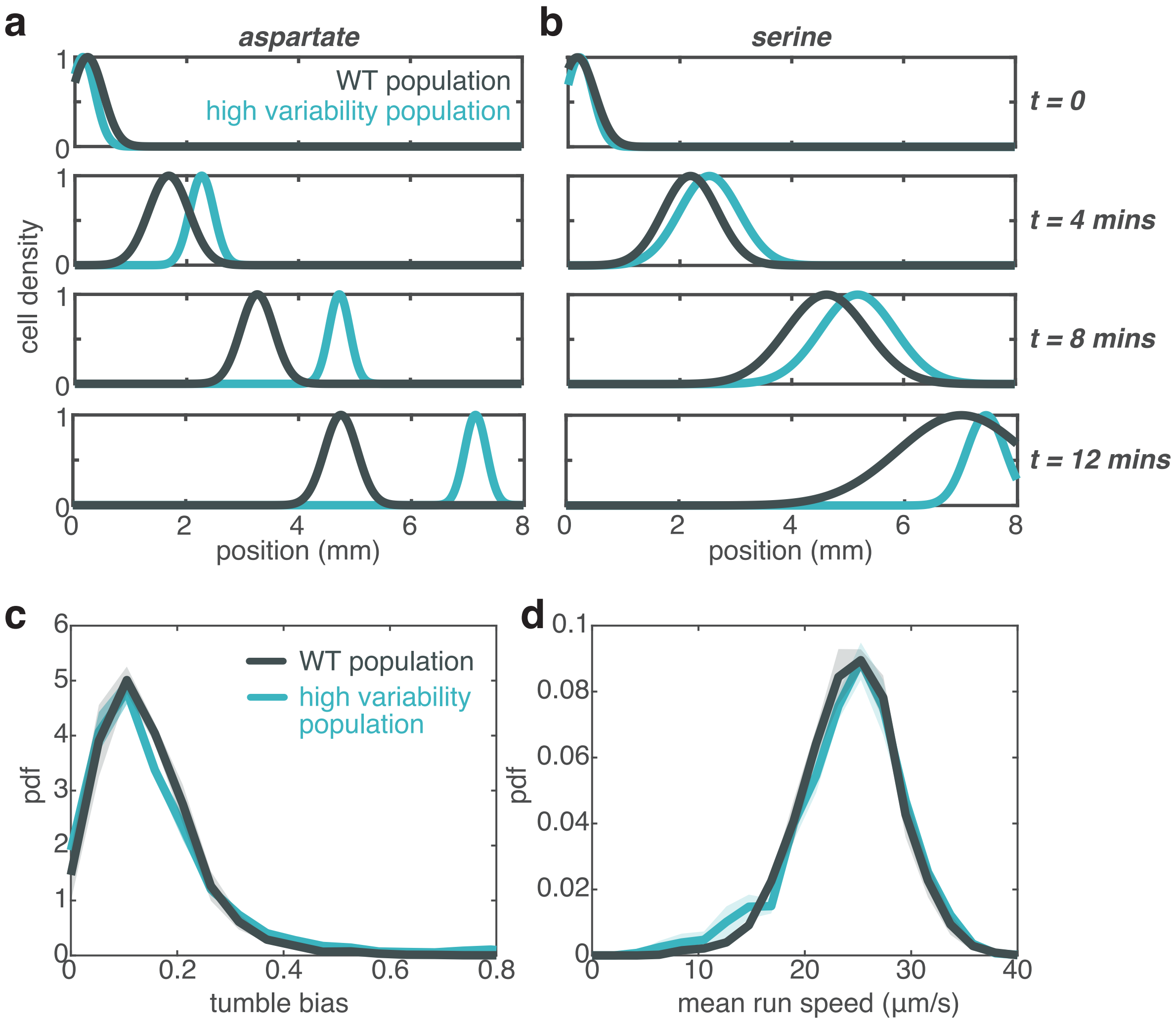


**EDF 6** Wave position, tumble bias, and run speed distributions of the WT and high variability populations. **(a)** Wave position of the high variability (left) and wild-type (WT) population (right) in a gradient of 100 $\mu$M L-aspartate. Cell density is extracted from the segmented wave images shown on figure 6b. Curves represent Gaussian function fits to the images. **(b)** Similar to panel a, but for cells in a gradient of 100 $\mu$M L-serine. **(c)** Tumble bias distribution of the high variability and wild-type populations in the absence of a chemical gradient are identical. Solid lines represent the mean of four independent biological replicates, with shaded areas indicating the standard error of the mean. **(d)** Same as panel a, but for the mean run speed of the high variability and wild-type populations.

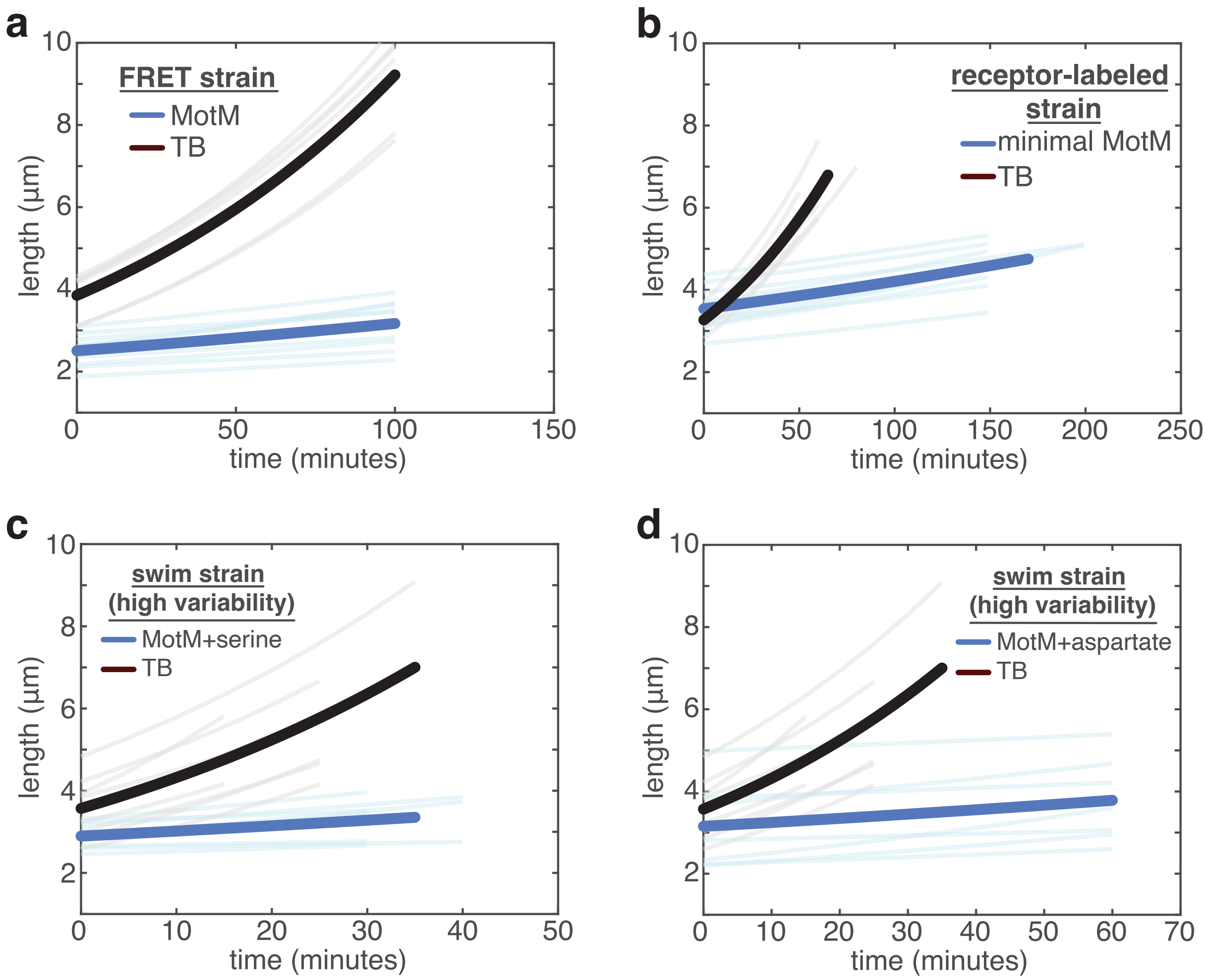


**EDF 7** Growth rate of the strains used in this study is negligible during experiments. **(a)** Cell elongation of the FRET strain (TSS2191/pSJAB106) as a function of time. Blue line represents the mean elongation rate of the stain in motility media (MotM), the buffer used for FRET experiments, while the black line represents the mean elongation rate of the same strain in TB, the rich media used to grow the strain. **(b)** Similar to panel a, but for the strain used in receptor-quantification experiments (TSS2155) in its measurement buffer, minimal motility media (minimal MotM). **(c)** Similar to panel a, but for the high variability strain used in swimming competition experiments (TSS2096/pTrc99A/pVS118) in MotM with 100 $\mu$M L-serine added. **(d)** Same as in panel c, but with 100 $\mu$M L-aspartate added to MotM.

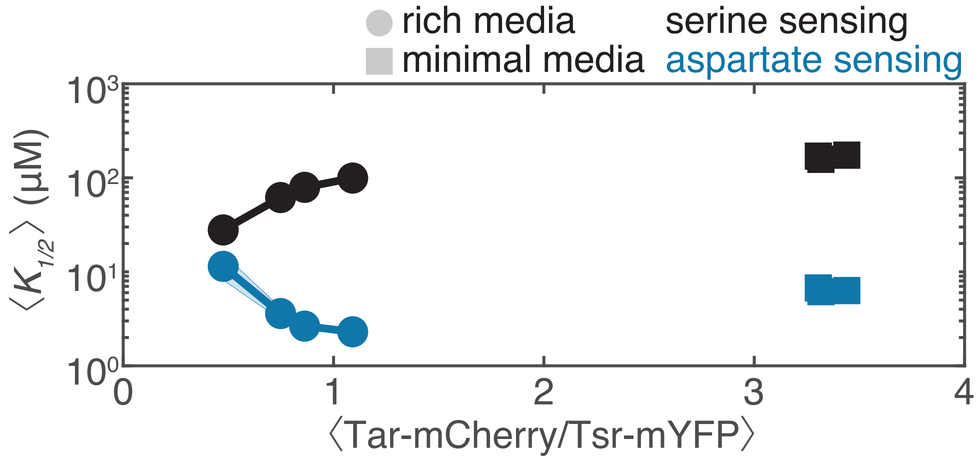


**EDF 8** Scaling of L-serine and L-aspartate ligand sensitivity $K_{1/2}$ as a function of receptor ratio. Scaling of mean L-serine (black) and L-aspartate (blue) $K_{1/2}$ as a function of mean receptor ratio for cells grown in rich (circles) and minimal media (squares). $K_{1/2}$ and receptor ratio were determined in separate experiments with cells grown under identical conditions. Shaded areas indicate 95% confidence intervals of the mean $K_{1/2}$ obtained via bootstrap resampling.

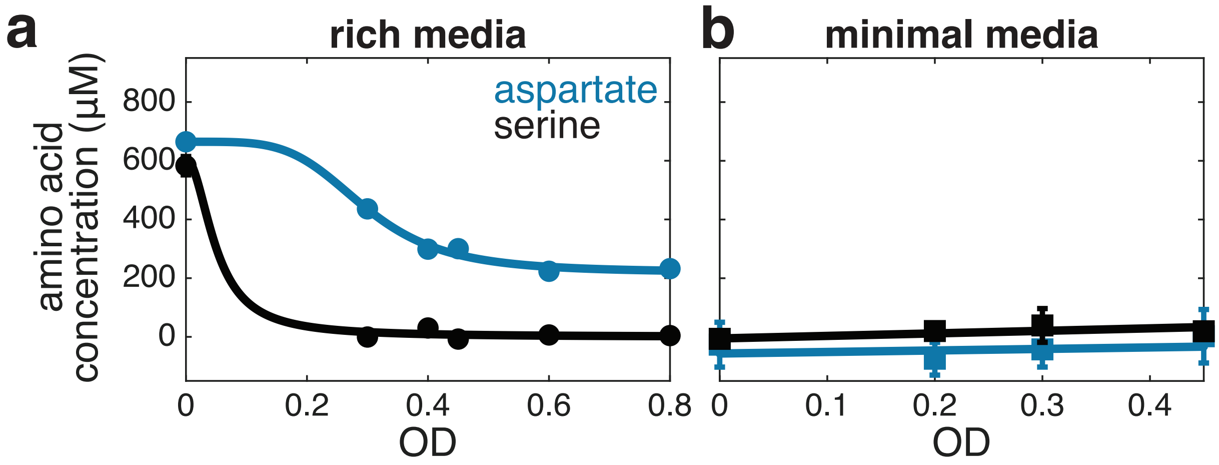


**EDF 9** Consumption of L-aspartate and L-serine in rich and minimal media. **(a)** Changes in L-aspartate and L-serine concentrations were measured in the supernatant of the FRET strain (TSS2191/pSJAB106) during growth in rich media as a function of cell density (OD), using a colorimetric assay (see Methods). Cells preferentially consume L-serine, which is depleted first, followed by consumption of L-aspartate. Supernatant samples were collected from the same growing culture at the indicated ODs, and both amino acid concentrations were quantified from each sample. Data points represent the mean of two biological replicates; error bars denote the standard error of the mean (SEM) and are typically on the order of the data point size. Solid lines represent sigmoidal fits to the data. **(b)** Same as in panel (a), but for cells grown in minimal media. Concentrations of both L-aspartate and L-serine remain below the detection limit throughout growth. Data points represent the mean of three biological replicates; error bars denote SEM. Solid lines represent robust linear fits to the data.

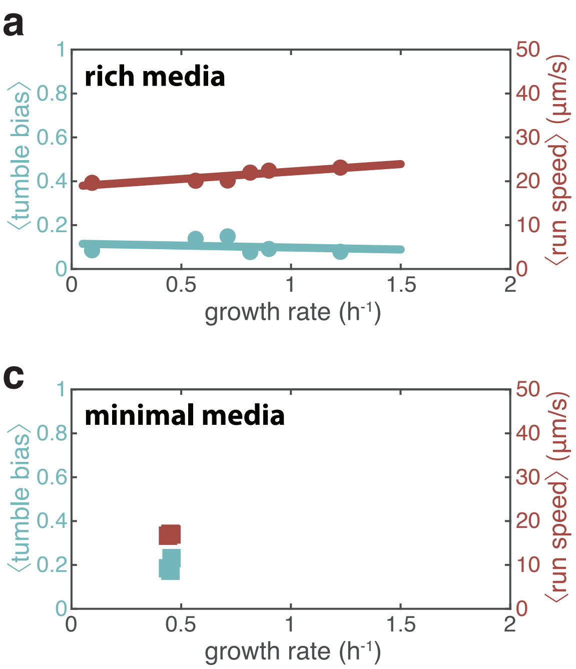


**EDF 10** Mean tumble bias and run speed as a function of growth rate and optical density. **(a)** Mean tumble bias and run speed of WT MG1655 as a function of growth rate for cells grown in rich media. Data from a single experiment. Lines are linear fits to the data. **(b)** Same data as panel (a) but plotted against the optical density (OD) of the culture. **(c)** Mean tumble bias and run speed of WT MG1655 as a function of growth rate for cells grown in minimal media. Data from a single experiment. **(d)** Same data as panel (c) but plotted against the optical density (OD) of the culture. Lines are linear fits to the data.

### References


[1] Keegstra, J.M., Kamino, K., Anquez, F., Lazova, M.D., Emonet, T., Shimizu, T.S.: Phenotypic diversity and temporal variability in a bacterial signaling network revealed by single-cell FRET. eLife **6**, 27455 (2017) https://doi.org/10.7554/eLife.27455

[2] Solari, J.: Spatial Organization of the Bacterial Cell: in Vivo Imaging Across Scales. Vrije Universiteit Amsterdam, Amsterdam, The Netherlands (2019). OCLC: 1088406919

[3] Mello, B.A., Tu, Y.: An allosteric model for heterogeneous receptor complexes: Understanding bacterial chemotaxis responses to multiple stimuli. Proceedings of the National Academy of Sciences **102**(48), 17354–17359 (2005) https://doi.org/10.1073/pnas.0506961102

[4] Cremer, J., Honda, T., Tang, Y., Wong-Ng, J., Vergassola, M., Hwa, T.: Chemotaxis as a navigation strategy to boost range expansion. Nature **575**(7784), 658–663 (2019) https://doi.org/10.1038/s41586-019-1733-y

[5] Qin, D., Xia, Y., Whitesides, G.M.: Soft lithography for micro- and nanoscale patterning. Nature Protocols **5**(3), 491–502 (2010) https://doi.org/10.1038/nprot.2009.234

[6] Kim, J.M., Garcia-Alcala, M., Balleza, E., Cluzel, P.: Stochastic transcriptional pulses orchestrate flagellar biosynthesis in *Escherichia coli*. Science Advances **6**(6), 0947 (2020) https://doi.org/10.1126/sciadv.aax0947

[7] Ronneberger, O., Fischer, P., Brox, T.: U-Net: Convolutional Networks for Biomedical Image Segmentation. arXiv. arXiv:1505.04597 [cs] (2015). http://arxiv.org/abs/1505.04597

[8] Kamino, K., Kadakia, N., Avgidis, F., Liu, Z.-X., Aoki, K., Shimizu, T., Emonet, T.: Optimal inference of molecular interaction dynamics in FRET microscopy. Proceedings of the National Academy of Sciences **120**(15), 2211807120 (2023) https://doi.org/10.1073/pnas.2211807120

[9] Phan, T.V., Mattingly, H.H., Vo, L., Marvin, J.S., Looger, L.L., Emonet, T.: Direct measurement of dynamic attractant gradients reveals breakdown of the Patlak–Keller–Segel chemotaxis model. Proceedings of the National Academy of Sciences **121**(3), 2309251121 (2024) https://doi.org/10.1073/pnas.2309251121

[10] Kamande, J.W., Wang, Y., Taylor, A.M.: Cloning SU8 silicon masters using epoxy resins to increase feature replicability and production for cell culture devices. Biomicrofluidics **9**(3), 036502 (2015) https://doi.org/10.1063/1.4922962

[11] Vo, L., Avgidis, F., Mattingly, H.H., Edmonds, K., Burger, I., Balasubramanian, R., Shimizu, T.S., Kazmierczak, B.I., Emonet, T.: Nongenetic adaptation by collective migration. Proceedings of the National Academy of Sciences **122**(8), 2423774122 (2025) https://doi.org/10.1073/pnas.2423774122

[12] Mattingly, H.H., Kamino, K., Machta, B.B., Emonet, T.: Escherichia coli chemotaxis is information limited. Nature Physics **17**(12), 1426–1431 (2021) https://doi.org/10.1038/s41567-021-01380-3
